# Combinatorial selective ER-phagy remodels the ER during neurogenesis

**DOI:** 10.1101/2023.06.26.546565

**Authors:** Melissa J. Hoyer, Cristina Capitanio, Ian R. Smith, Julia C. Paoli, Anna Bieber, Yizhi Jiang, Joao A. Paulo, Miguel A. Gonzalez-Lozano, Wolfgang Baumeister, Florian Wilfling, Brenda A. Schulman, J. Wade Harper

**Affiliations:** Department of Cell Biology, Harvard Medical School, Boston MA 02115; Aligning Science Across Parkinson’s (ASAP) Collaborative Research Network, Chevy Chase, MD 20815, USA; Department of Molecular Machines and Signaling, Max Planck Institute of Biochemistry, 82152 Martinsried, Germany; Department of Molecular Structural Biology, Max Planck Institute of Biochemistry, 82152 Martinsried, Germany; Mechanisms of Cellular Quality Control, Max Planck Institute of Biophysics, 60438 Frankfurt a. M., Germany

**Author notes:** equal contribution (Cristina Capitanio, Ian R. Smith).

## Abstract

The endoplasmic reticulum (ER) employs a diverse proteome landscape to orchestrate many cellular functions ranging from protein and lipid synthesis to calcium ion flux and inter-organelle communication. A case in point concerns the process of neurogenesis: a refined tubular ER network is assembled via ER shaping proteins into the newly formed neuronal projections to create highly polarized dendrites and axons. Previous studies have suggested a role for autophagy in ER remodeling, as autophagy-deficient neurons *in vivo* display axonal ER accumulation within synaptic boutons, and the membrane-embedded ER-phagy receptor FAM134B has been genetically linked with human sensory and autonomic neuropathy. However, our understanding of the mechanisms underlying selective removal of ER and the role of individual ER-phagy receptors is limited. Here, we combine a genetically tractable induced neuron (iNeuron) system for monitoring ER remodeling during *in vitro* differentiation with proteomic and computational tools to create a quantitative landscape of ER proteome remodeling via selective autophagy. Through analysis of single and combinatorial ER-phagy receptor mutants, we delineate the extent to which each receptor contributes to both magnitude and selectivity of ER protein clearance. We define specific subsets of ER membrane or lumenal proteins as preferred clients for distinct receptors. Using spatial sensors and flux reporters, we demonstrate receptor-specific autophagic capture of ER in axons, and directly visualize tubular ER membranes within autophagosomes in neuronal projections by cryo-electron tomography. This molecular inventory of ER proteome remodeling and versatile genetic toolkit provides a quantitative framework for understanding contributions of individual ER-phagy receptors for reshaping ER during cell state transitions.

## MAIN

The ER network is shaped by proteins that promote tubule and sheet-like membrane structures, which in turn tailors ER function in a cell type specific manner to optimize protein secretion, calcium storage, lipid homestasis, and inter-organelle contacts^1-3,4^. ER-phagy represents a central mechanism through which ER can be remodeled, or superfluous ER proteins or lipids recycled^5,6^. Several membrane-embedded ER proteins have been implicated in ER remodeling via ER-phagy in different cellular contexts with varying degrees of evidence^5,7^. Membrane bound ER-phagy receptors include single pass transmembrane (TM) segment containing proteins TEX264, CCPG1, SEC62 and reticulon-like hairpin domain (RHD) containing FAM134A, B, C, (also called RETREG2, 1, 3, respectively), Atlastin (ATL2), and RTN3L^8-16^. RHDs are thought to reside in the outer leaflet of the ER membrane to induce curvature^17-19^. All identified ER-phagy receptors contain cytosolic LC3-interaction region (LIR) motifs that bind to ATG8 proteins such as MAP1LC3B (also called LC3B) on the phagophore to promote ER capture^5^. Mechanistically, the FAM134 class of receptors are thought to cluster through their hairpin RHDs into highly curved nanoscale membrane domains that recruit autophagy machinery to emerging ER membrane “buds”, thereby nucleating phagophore formation^5,6,20-22^. Phagophore expansion and ultimate autophagosome closure around ER is thought to be coupled to scission of ER membrane at the bud neck, although the underlying mechanism is yet to be determined.

Given the complexity of ER-phagy receptors and the fact that most studies have involved either nutrient stress or receptor overexpression models, central unanswered questions in the field include when, where and how individual receptors are used to remodel ER during physiological changes in cell state. In addition, there is limited assessment as to the extent to which individual ER proteins are actively cleared as ER-phagy “cargos” in unique cell states. While ER protein accumulation has been observed in synaptic boutons of mouse primary hippocampal neurons from autophagy-deficient ATG5^-/-^ mice^23^, it was suggested that ER proteome accumulation upon autophagy inhibition in neurons reflects non-selective autophagy rather than selective ER-phagy^23^. An understanding of ER-phagy is further confounded by ER membranes both serving as a source of phospholipids for autophagosome expansion^24^ *and* as being captured as cargo within a fully formed autophagosome via selective ER-phagy, as visualized by electron microscopy^10,11^. However, critical work has revealed that the process of lipid transfer from the ER to the growing autophagosome occurs *without* incorporation of ER proteins into the phagophore membrane itself^24,25^. Thus, the process of ER-phagy receptor facilitated ER protein clearance is functionally and mechanistically distinct from the use of ER membranes as a source for phospholipids in phagophore expansion.

Here, we employ an in vitro neurogenesis system that recapitulates central autophagy-dependent features of ER remodeling^26^ to directly examine the role of ER-phagy receptors in this process. We identify redundant and selective ER-phagic cargo for individual receptors, demonstrate a role for multiple ER-phagy receptors in eliminating axonal ER, directly visualize ER-phagy receptors trafficking in autophagosomes in axons, and visualize tubular ER membranes captured within autophagosomes in neuronal projections via cryo-electron tomography (cryo-ET). Unlike proteins such as neurogenesis or pluripotency factors which display large magnitude changes in their abundance during conversion of ES cells to induced neurons (iNeurons), we find that ER protein remodeling by autophagy represents a continuum of comparatively small changes in abundance of individual ER-resident proteins. This feature necessitated the development of a quantitative proteomic framework capable of measuring and classifying abundance changes across the ER proteome and in the context of an allelic series of ER-phagy receptor mutants. We find that FAM134 family members play a dominant and largely redundant role in remodeling ER membrane proteins during neurogenesis, while CCPG1 is primarily responsible for autophagic turnover of a cohort of lumenal ER proteins, thereby defining an underlying specificity for ER proteome remodeling. These data provide a proteomic landscape for ER remodeling in iNeurons and an experimental framework for elucidating how changes in cell state control the ER proteome via selective autophagy.

### Landscape of ER remodeling by autophagy during in vitro neurogenesis

The ER proteome is composed of ∼350 proteins in four broad categories^27^: 1) membrane spanning proteins harboring one to as many as 14 TMs, 2) ER-associated proteins which are cytosolic but interact with ER-membrane, 3) lumenal ER proteins, and 4) ER tubule and sheet shaping proteins (**Extended Data Fig. 1a and Supplementary Table 1**). To examine alterations in ER protein abundance, we initially mined total proteome abundance measurements from our previously described system wherein NGN2 induction in human ES cells drives neurogenesis with >95% efficiency^26^ (**Fig. 1a**). During a 12-day iNeuron differentiation, a diverse cohort of ER proteins within multiple functional categories increase or decrease in abundance (**Fig. 1a-b, Extended Data Fig. 1b, Supplementary Table 2**)^26^. Proteins undergoing the largest increase in abundance include enzymes involved in protein folding (e.g. FKBP9, CRELD1), ion regulation (e.g. RCN1, TMEM38B), and collagen modification (e.g. COL4A2), while other collagen modifying proteins PXDN and P3H4 decreased in abundance (**Fig. 1a**). Among ER shaping proteins, RHD containing proteins RTN1, RTN4, and REEP2 all displayed the largest increase in abundance, with greater than 1.4-fold increases from day 0 levels. which is represented by Log_2_ (fold change day 12 versus day 0) greater than 0.5, or more simply shown as Log_2_FC>0.5 (**Fig. 1b, Extended Data Fig. 1b**). These results are consistent with the formation of extensive ER tubule networks within neuronal projections which has been previously characterized^28^. Indeed, immunofluorescence of ER tubular shaping protein RTN4 revealed extensive RTN4-positive projections in iNeurons, while the ER sheet protein CKAP4 (also called CLIMP63) was largely confined to the cell body (soma) (**Extended Data Fig. 1c**).

**Fig. 1.**
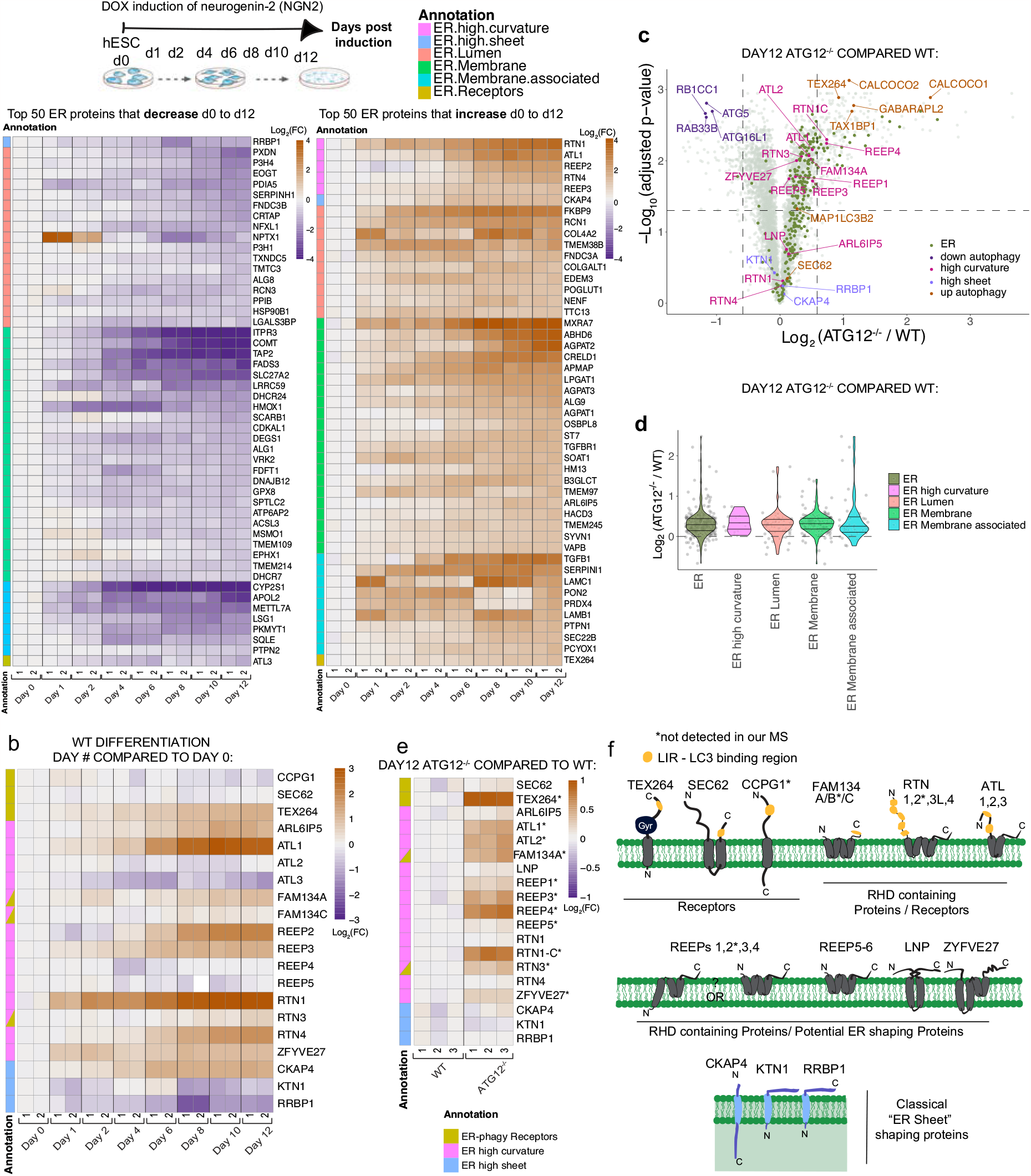
Landscape of ER remodeling via autophagy during hESC differentiation to iNeurons in vitro. **a**, Changes in abundance of the most highly remodeled ER proteins during conversion of WT hESCs to iNeurons are shown in heatmaps (Log_2_ Fold Change (FC) at indicated day of differentiation relative to hESCs). The top 50 proteins that either decrease or increase in abundance are shown; see **Extended Data Fig. 1b** for full heatmap. Data are from our previous analysis of iNeuron differentiation. Annotations depicting type of ER protein are indicated by the relevant colors. **b** Heat map (Log_2_FC) of ER shaping proteins specifically in differentiating iNeurons **c**, Volcano plot [-Log_10_ (adjusted p-value) versus Log_2_FC (ATG12^-/-^/WT)] of day 12 WT and ATG12^-/-^ iNeuron total proteomes displaying accumulation of autophagy-related and ER proteins (green dots) as a cohort. Each dot represents the average of triplicate TMT measurements. **d**, Violin plots for individual classes of ER proteins showing the relative increases in abundance in ATG12^-/-^ day 12 iNeurons compared with WT iNeurons. Each dot represents the average of triplicate TMT measurements. **e** Heat map (Log_2_FC) of ER shaping proteins specifically in day 12 WT versus ATG12^-/-^ iNeurons. Asterisk after gene name indicates significant changes in abundance: *, adjusted p<0.05. **f**, Topology of ER shaping proteins and ER-phagy receptors within the ER membrane. Annotation color scheme for individual classes of ER proteins in **e** also applies to **b**.

We next compared wildtype (WT) and ATG12^-/-^ day 12 iNeurons using Tandem Mass Tagging (TMT) proteomics (**Fig. 1c, Supplementary Table 3**). ATG12 is a canonical autophagy protein and is conjugated to ATG5 to support lipidation of ATG8 proteins. Consistent with previous results^26^, several ubiquitin binding autophagy receptors (CALCOCO1, CALCOCO2, TAX1BP1) and the ATG8 protein GABARAPL2 accumulated in ATG12-deficient iNeurons, as did ER-phagy receptors TEX264 and FAM134A (greater than 1.4-fold change, Log_2_ FC> 0.49) (**Fig. 1c**). These results are consistent with the expected reduction in autophagic clearance. Moreover, a cohort of ER proteins displayed increased abundance, as indicated by the rightward skew distribution in volcano plots of Log_2_ fold change (FC) values for ATG12^-/-^/WT proteomes. Similarly, violin plots revealed an overall increase in ER protein abundance, which showed a mean Log_2_FC of 0.33, representing a mean 1.26-fold increase across the entire cohort of ER proteins (**Fig. 1c-d, Extended Data Fig. 1d**). Strikingly, RHD proteins accumulate to the greatest degree (including REEP1-4, RTN1), while classical ER sheet proteins CKAP4 and RRBP1 were largely unchanged (**Fig. 1d-f**). Alterations in protein abundance for TEX264, REEP5 and CKAP4 were verified by immunoblotting, as was increased abundance of FAM134C (not detected by proteomics in this experiment) (**Extended Data Fig. 1e,f**). Mapping the landscape of ER protein accumulation in ATG12 deletion iNeurons (Log_2_ FC from WT) revealed that, beyond ER curvature shaping proteins, specific ER proteins assigned to several other structural or functional categories accumulate during differentiation in the absence of autophagy, including lumenal and transmembrane segment-containing biosynthetic or metabolic proteins (**Extended Data Fig. 1a**). Importantly, the extent of differentiation efficiency of ATG12^-/-^ iNeurons was equivalent to WT iNeurons as assessed by the induction of several neuronal differentiation markers and loss of pluripotency factors (**Extended Data Fig. 2a, Supplementary Table 4**). Moreover, the viability of ATG12^-/-^ iNeurons was equivalent with WT iNeurons (**Extended Data Fig. 2b-c**), which is consistent with the similar differentiation efficiency results. We also directly examined the possibility that loss of ATG12 promotes ER stress in day 12 iNeurons, but detected no increase in the ER stress response markers ATF4 (protein level expression) or *XBP-1* (mRNA splicing) when compared WT cells, while tunicamycin as a positive control induced both ATF4 expression and *XBP-1* splicing (**Extended Data Fig. 2d,e**). Taken together, we conclude that differentiation of ES cells to iNeurons in the absence of ATG12 is associated with alterations in the abundance of the ER proteome.

### Aberrant axonal ER accumulation during neurogenesis without autophagy

We next examined ER morphology in WT or ATG12^-/-^ day 20 iNeurons using α-Calnexin or α-RTN4 as general or tubule-enriched markers for ER, respectively. Day 20 iNeurons were employed here as they lie flatter and projections are therefore more easily imaged than day 12 iNeurons. We observed ER-positive accumulations that dilated the projections in autophagy-deficient cells. These ER accumulations were both larger and more numerous than those seen in WT iNeurons (**Fig. 2a,b, Extended Data Fig. 3a**). Immunofluorescent staining with α-NEFH (high molecular weight neurofilament-H) verified that the projections were present within axons (**Fig 2b**). Interestingly, α-NEFH-positive filaments formed a “cage-like” structure with multiple filaments encasing the ER (**Fig 2b, inset**). The mean area of ER accumulations dilating the axons in ATG12^-/-^ iNeurons (in α-NEFH positive axon regions) was 12.2 micron^2^, while in WT iNeurons these were less abundant and consistently smaller (mean area 6.3 micron^2^) (**Fig. 2b, Extended Data Fig. 3a**). To examine these structures at higher resolution, we employed scanning transmission electron microscopy (TEM) (**Extended Data Fig. 3b,c**). As seen with light microscopy, dilated structures within WT processes were more rarely observed by TEM. Membrane and organelles were present in the WT axonal dilated structures that did form, and in one example, a WT bulbous structure was rich in ER. Analogous, but larger dilated ER-containing structures were frequently found in TEM images of ATG12^-/-^ iNeurons (**Extended Data Fig. 3b,c**). The ER networks in these examples forms around or adjacent to the continuous microtubule tracks. These dilated axonal regions filled with ER are reminiscent of previously observed axon boutons within mouse neurons lacking *Atg5*^23^.

**Fig. 2.**
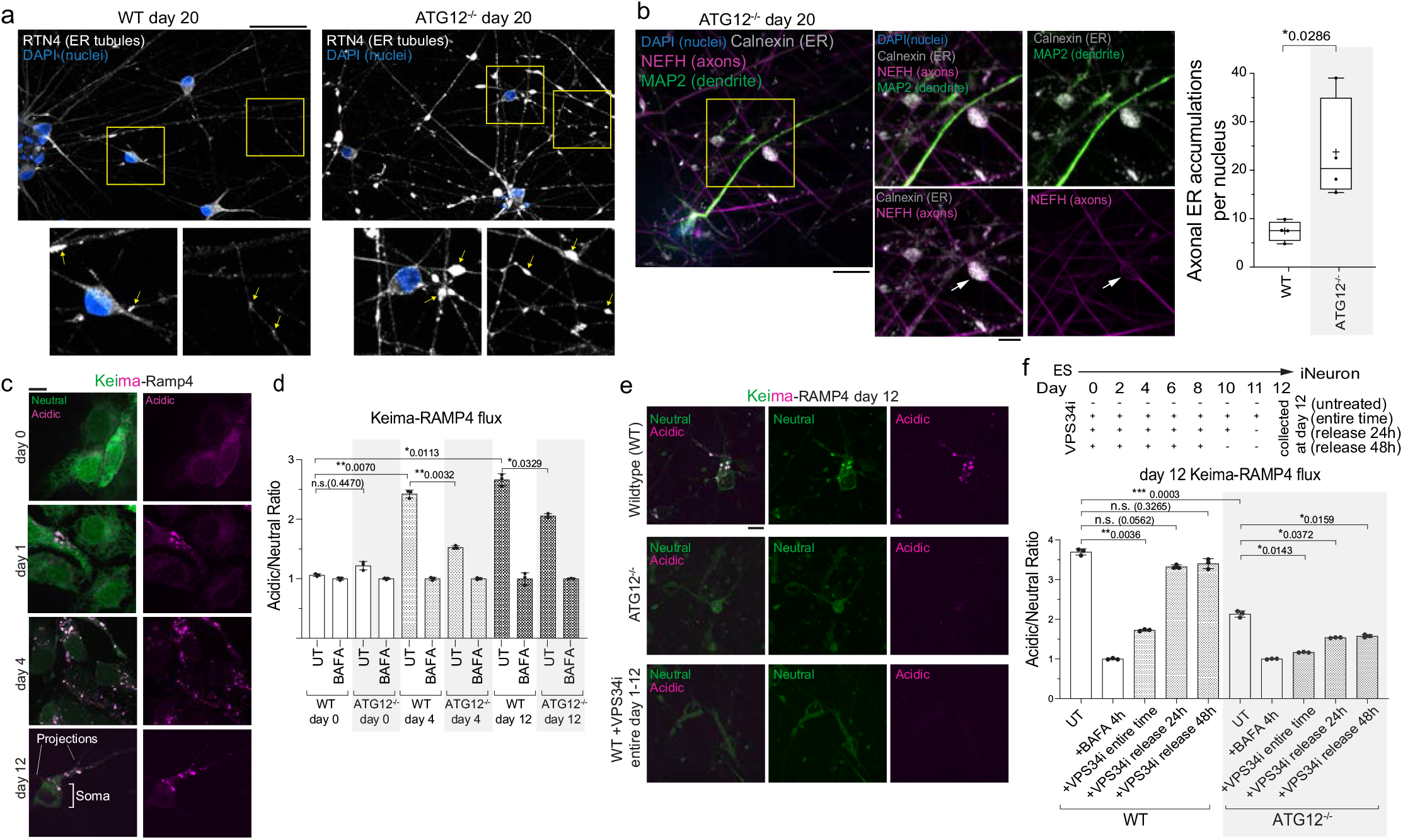
ER-phagic flux in iNeurons. **a**, WT or ATG12^-/-^ day 20 iNeurons immunostained with ER-tubule marker α-RTN4 (white) and with DAPI (nuclei, blue). Scale bar, 50 microns. **b**, Enlarged ER-positive structures in ATG12^-/-^ day 20 iNeurons revealed by immunostaining with α-Calnexin, ER (white); α-MAP2, dendrites (green); α-NEFH, axons (magenta); and DAPI, nuclei (blue). Scale bar, 10 microns. Right panel, min-to-max box-and-whiskers plot representing mean number of axonal ER accumulations per nucleus. Line marks median and + labels the mean. Points represent mean values from four independent differentiations. *, p<0.05, Mann-Whitney test. **c**, hESCs expressing Keima-RAMP4 were differentiated to iNeurons. Keima was imaged at day 0, 1, 4 and 12. Scale bar, 5 microns **d**, WT or ATG12^-/-^ Keima-RAMP4 flux was measured by flow cytometry at day 0, 4 and 12 of differentiation. The ratio of acidic to neutral Keima fluorescence was normalized to samples treated with BAFA (100 nM, 4 h). **e**, Reduced Keima-RAMP4 flux in ATG12^-/-^ iNeurons or upon VPS34 inhibitor, VPS34i (1 μM) treatment was measured as in **c**. Scale bar, 10 microns for **c** and **e. f**, WT or ATG12^-/-^ hESCs differentiated with or without VPS34i as indicated in the scheme. In some conditions, VPS34i was washed out at time indicated (24 or 48 h) prior to analysis. In **d** and **f**, each point represents one of three biological triplicate measurements. Error bars represent SD. *, p<0.05; **, p<0.01; ***, p<0.001; n.s., not significant; Brown-Forsythe and Welch One-way ANOVA and Dunnett’s T3 multiple comparisons test.

### ER-phagic flux during differentiation and in iNeurons

To characterize the temporal properties of ER accumulation in ATG12^-/-^ iNeurons, we next measured ER protein clearance to the lysosome (ER-phagic flux) at different stages of differentiation and in post-differentiated “established” iNeurons. We linked pH sensitive Keima to pan-ER (Keima-RAMP4, a reporter widely used in the ER-phagy field) (**Fig. 2c,d; Extended Data Fig. 3d**) or to ER tubules (Keima-REEP5) (**Extended Data Fig. 3e,f**) and compared nonacidified Keima-ER throughout the ER network to acidified Keima-ER that had reached the low pH environment of lysosomes^10,14,29^. Neither reporter underwent significant flux to lysosomes in day 0 ES cells, consistent with our previous data showing ER proteins do not accumulate in ATG12^-/-^ ES cells^26^. However, during differentiation, we observed an increase in acidic Keima signal (increased acidic/neutral ratio as defined in **METHODS**) for both ER reporters, with acidified puncta representing ER in lysosomes located primarily in the soma (**Fig. 2c; Extended Data Fig. 3d,e**). Parallel experiments using flow cytometry quantified the amount of ER flux to lysosomes upon differentiation using both reporters (**Fig. 2d; Extended Data Fig. 3f,g**). Acidic signal was normalized to cells treated with Bafilomycin A (BAFA, 4h), which pharmacologically inhibits lysosomal acidification. This ER flux was substantially reduced in cells lacking ATG12, and residual flux was further eliminated by addition of the small molecule VPS34 PI3 kinase inhibitor SAR405 (VPS34i) throughout the differentiation time course, which blocks phagophore initiation (**Fig. 2e,f**). Detectable flux in ATG12^-/-^ cells is consistent with the previous finding that loss of the ATG8 conjugation system does not fully block autophagosome formation^30^. Due to the long half-life of Keima in lysosomes^31^, detectable stable Keima within lysosomes over multiple days of differentiation was expected. Release from continuous VPS34 inhibition one or two days prior (at day 10 or 11) to neuron collection (at day 12) resulted in increased Keima flux to lysosomes that was comparable to flux in untreated cells; this increase was absent in cells lacking ATG12 (**Fig. 2f**). Finally, we examined whether ER-phagic flux was ongoing in established iNeurons. Keima flux measured in later stage day 20 neurons was reduced by adding VPS34i at day 15 of differentiation, as compared with untreated cells (**Extended Data Fig. 3g**). These results indicate that ER fluxes to lysosomes during differentiation in a process that requires canonical autophagy, and that autophagic ER flux is ongoing in established iNeurons.

### ER-phagy receptor capture by autophagosomes in axons and somata

Our findings left the localization of ER-phagy an apparent paradox: we observed acidic Keima-RAMP4 puncta within the soma (**Extended Data Fig. 3d**) but also detected dramatic ER accumulation within axons of ATG12^-/-^ iNeurons (**Fig. 2a,b, Extended Data Fig. 3a-d**). It is well known that autophagosomes originate in axons, fuse with lysosomes to form autophagolysosomes that acidify during retrograde trafficking en-route to the soma^32-35^. Thus, acidic Keima-RAMP4-positive puncta in the soma could reflect ER-phagy occurring locally in the soma or, alternatively, axonal ER-phagic capture within autophagosomes followed by retrograde transport to the soma where there is fusion with acidic lysosomes.

To examine spatial aspects of ER-receptor capture, we expressed TEX264-GFP or FAM134C-GFP in iNeurons (**Extended Data Fig. 4a-b**). Previous studies in non-neuronal cell lines demonstrated that both TEX264 and FAM134 proteins can localize broadly throughout the ER network and can form co-incident puncta that become engulfed by autophagosomes^8,10^. We observed TEX264-GFP punctate structures (indicated by arrowheads) both in projections and in the soma (at day 4 of differentiation) that were rarely detected: 1) when TEX264’s LIR motif was mutated (F273A mutant), 2) in cells lacking ATG12, or 3) in cells treated with VPS34i (**Extended Data Fig. 4a,c**). These results suggest that ER-phagy receptor puncta formation in iNeurons was likely due to active ER-phagy as described previously in other cell systems that used starvation as a trigger for ER-phagy^8,10^.

To further verify TEX264-GFP puncta in autophagic structures, we co-expressed mCherry-LC3B (mCh-LC3B). Co-staining with α-NEFH in fixed cells verified co-incidence of mCh-LC3B and TEX264-GFP in axons (**Extended Data Fig. 4d**). We took advantage of the highly polarized and defined axons in live day 30 iNeurons to track the movement of mCh-LC3B/TEX264-GFP-positive puncta. Numerous GFP-TEX264 puncta trafficked with mCh-LC3-positive structures (**Fig. 3a,b, Movie 1**). In each axon, autophagosomes enriched in TEX264 moved primarily unidirectionally (this predominant movement in one direction on the track is defined here as forward), but we also recorded stops and some backwards movements on these tracks (**Fig. 3c**). The median forward speed was 0.297 micron per second (**Fig. 3c**), which is similar to speeds previously reported for autophagosomes undergoing microtubule-dependent trafficking in axons of mouse primary neurons^36^. Similarly, dynamic FAM134C-GFP positive structures, trafficking with mCh-LC3B puncta, were also observed in day 30 iNeurons (**Fig. 3d, Movie 2**), indicating that multiple ER-phagy receptors may be operating within projections.

**Fig. 3.**
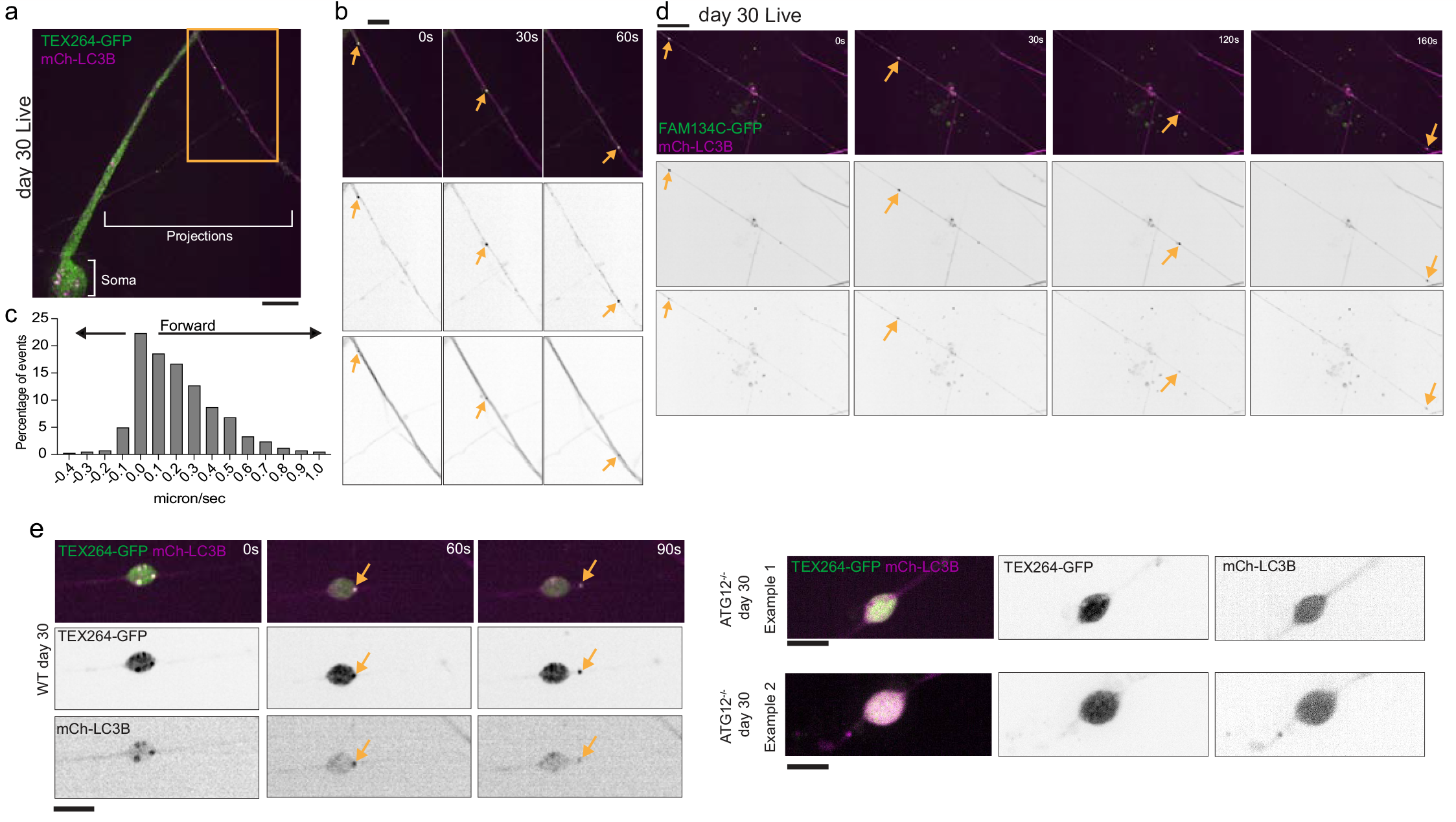
Axonal trafficking of TEX264-GFP and FAM134C-GFP-containing autophagosomes via live-cell imaging. **a**,**b**, TEX264-GFP (green) and mCh-LC3B (magenta) day 30 iNeurons imaged live (panel **a**). Inset in panel **b** shows positions of mCh-LC3B/TEX264-GFP-positive puncta trafficking within an axon. Arrowheads indicate puncta positions over two indicated time sequences. Scale bars; 10 microns and 5 microns for **a** and **b**, respectively. **c**, Rate of TEX264-GFP/mCh-LC3B-positive puncta movements (n=429), and the percentage of events at indicated speeds are binned in a histogram (events from three replicate differentiation experiments). **d**, As in **b**, but for FAM134C-GFP/mCh-LC3B-positive puncta. **e**, TEX264-GFP/mCh-LC3B-positive puncta are in dilated regions of WT iNeuron axons and traffic away (left panels), but puncta are not detected in ATG12^-/-^ iNeurons (right panels). Scale bars, 10 microns.

We next examined the question of whether ER-rich axonal dilations might be sites of ER-phagic capture. Indeed, live-cell imaging revealed TEX264-GFP/mCh-LC3B-positive puncta emerging from small axonal dilations in WT neurons (**Fig. 3e, Movie 3**). In contrast, while TEX264-GFP was present in regions with dilated axonal ER in ATG12^-/-^ iNeurons, TEX264-GFP/mCh-LC3B-positive puncta were not observed (**Fig. 3e**). These data suggest a role for ER-phagy receptor-dependent clearance of ER in axonal processes.

### Visualization of tubular ER within autophagosomes in neuronal projections by cryo-ET

TEX264-GFP trafficking co-incident with mCh-LC3B led us to ask whether ER-containing autophagosomes could be visualized along microtubules in neuronal projections by cryo-electron tomography (cryo-ET). We established a workflow in which iNeurons grown on EM grids were plunge-frozen at day 18, followed by cryo-fluorescence microscopy and cryo-ET of thin neuronal projections (**Fig. 4a;** see METHODS for details). We used an unbiased approach to survey axonal projections for autophagic structures, which we identified directly in TEM images. We then correlated autophagosome positions in TEM images with cryo-fluorescence data to evaluate coincidence with fluorescence signal. Finally, neural network-based segmentation revealed the cargo and cellular surroundings of the captured autophagosomes. Our final dataset contained 37 autophagosomes captured *in situ* in axonal projections (**Fig. 4b-h, Extended Data Fig. 5a-f**, see METHODS). Many autophagic structures (24/37) were proximal to microtubules within the axon, as would be expected during trafficking to the soma (**Fig. 4c-h, Extended Data Fig. 5d,f**). Interestingly, tubular ER is clearly present as cargo inside 21 of the 37 autophagosomes analyzed (**Fig. 4c-h, Extended Data Fig. 5e,f, Movie 4**), and regions with GFP signal were co-incident with ER tubule-containing autophagic structures (**Fig. 4b,d,g, Extended Data Fig. 5g-n**). In particular, 5/21 ER tubule-containing autophagic structures coincided with TEX264-GFP signal and 4/5 of these were also linked to microtubules. Absence of GFP-TEX264 signal in a subset of ER-containing autophagosomes may be due to low expression level of lentiviral-transduced TEX264-GFP in individual iNeurons or due to selective capture by ER-phagy receptors other than TEX264. Note that while TEX264-GFP displayed punctate and distinct signal, mCh-LC3B cryo-fluorescence signal appeared diffuse and could not be reliably used for correlation (**Extended Data Fig. 5m**; see METHODS). Notably, none of the autophagosomes which did not contain tubular ER cargo coincided with TEX264-GFP signal (**Fig. 4b)**. Taken together, these data confirm capture of TEX264-GFP-positive ER by autophagy within axonal projections and is consistent with the occurrence of selective ER-phagy in iNeurons.

**Fig. 4.**
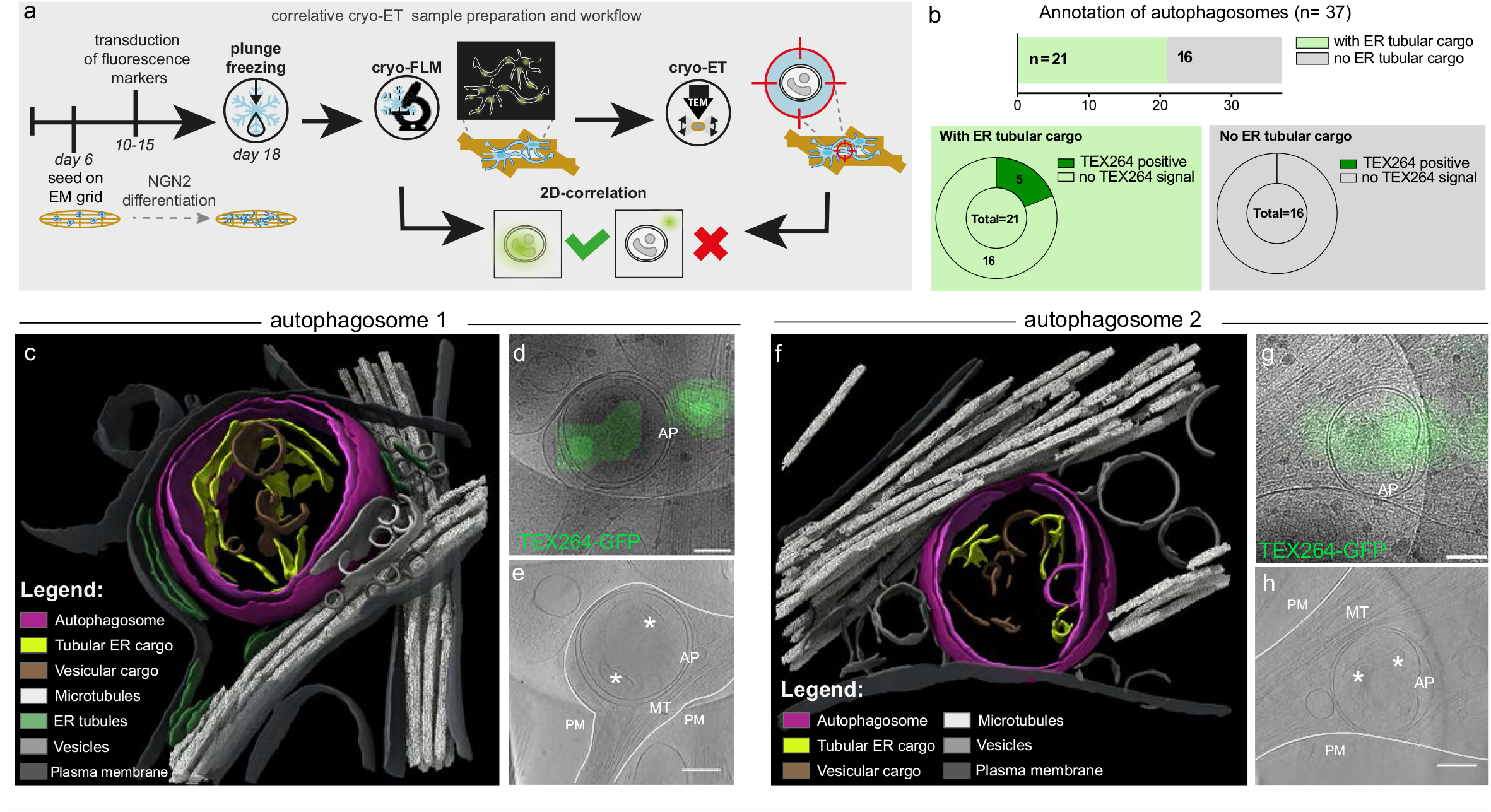
Observation of tubular ER within autophagosomes in neuronal projections by correlative cryo-ET. **a**, Experimental strategy used to capture autophagosomes in the projections of iNeurons. Induced pluripotent stem cells (iPSCs) are differentiated on EM grids and transduced with fluorescence markers before plunge-freezing at day 18. After imaging the sample by cryo-fluorescence microscopy (cryo-FLM), autophagosomes in the neuronal projections are identified in TEM images based on morphological features and captured by cryo-electron tomography (cryo-ET). 2D-correlation of TEM images with previously acquired fluorescence data shows whether autophagosomes correlate or not with fluorescence markers, such as TEX264-GFP. **b**, Cargo and TEX264-GFP correlation analysis of captured autophagosomes. The barplot on top shows the number of autophagosomes in which ER tubular cargo is present (green, n= 21) or not (gray, n=16). The pie charts show the number of structures corresponding to TEX264-GFP signal in each category. **c-h**, Examples of TEX264-GFP positive autophagosomes with tubular ER cargo captured *in situ* by cryo-ET. **c**,**f**, 3D segmentations reveal double membrane autophagosomes (magenta) containing ER tubules as cargo (yellow) and close to microtubules (white). The tubular ER cargo of autophagosome 1 (c) exhibits a similar morphology as the adjacent cytosolic ER (green). For the full tomogram movie of autophagosome 1, see **Supplemental Movie 4**. For full segmentation of ER tubules, see **Extended Data Fig. 5e. d,g**, Zoomed-in 11500X TEM images corresponding to autophagosomes 1 and 2 overlayed with TEX264-GFP cryo-fluorescence signal. For a complete view of fluorescence overlays, see **Extended Data Fig. 5k,m. e,h**, Tomogram slices of autophagosomes 1 and 2, denoised with cryo-CARE. White lines indicate the plasma membrane (PM) of the neuronal projections containing the autophagosomes. Asterisks indicate the tubular ER cargo visible in these slices. AP, autophagosome; MT, microtubules. All scale bars, 200 nm.

### A genetic toolkit for ER-phagy receptor analysis in iNeurons

To systematically explore contributions of individual ER-phagy receptors to ER remodeling during iNeuron differentiation^37^, we used gene editing to first create single knock-out hESCs for FAM134A, FAM134B, FAM134C, TEX264, or CCPG1, which were confirmed by sequence analysis and immunoblotting of hESC extracts (**Fig. 5a, Extended Data Fig. 6a,b**). Multiple ER-phagy receptors accumulate in ATG12^-/-^ cells during differentiation and previous studies in non-neuronal cells indicate partial redundancy among ER-phagy receptors^11^, so we also sequentially edited FAM134C^-/-^ cells to create double, triple, quadruple, and penta receptor knockout lines: FAM134A/C^-/-^ (DKO), FAM134A/B/C^-/-^ (TKO), FAM134A/B/C/TEX264^-/-^ (QKO) and FAM134A/B/C/TEX264/CCPG1^-/-^ (PKO) (**Fig. 5b, Extended Data Fig. 6c,d**). Sequential deletion of ER-phagy receptors was verified by sequence analysis and immunoblotting; QKO and PKO mutants displayed normal karyotypes (**Extended Data Fig. 6, c,d,e**). As with ATG12^-/-^ and WT iNeurons, PKO iNeurons differentiated efficiently, displayed viability parameters equivalent to WT iNeurons, and displayed no evidence of ER stress as assessed by either ATF4 or *XBP1s* induction (**Extended Data Fig. 2a-e, Supplementary Table 4**). Each mutant cell line was reconstituted with Keima-RAMP4 to measure ER-phagic flux to the lysosome (**Fig. 5a,b**).

**Fig. 5.**
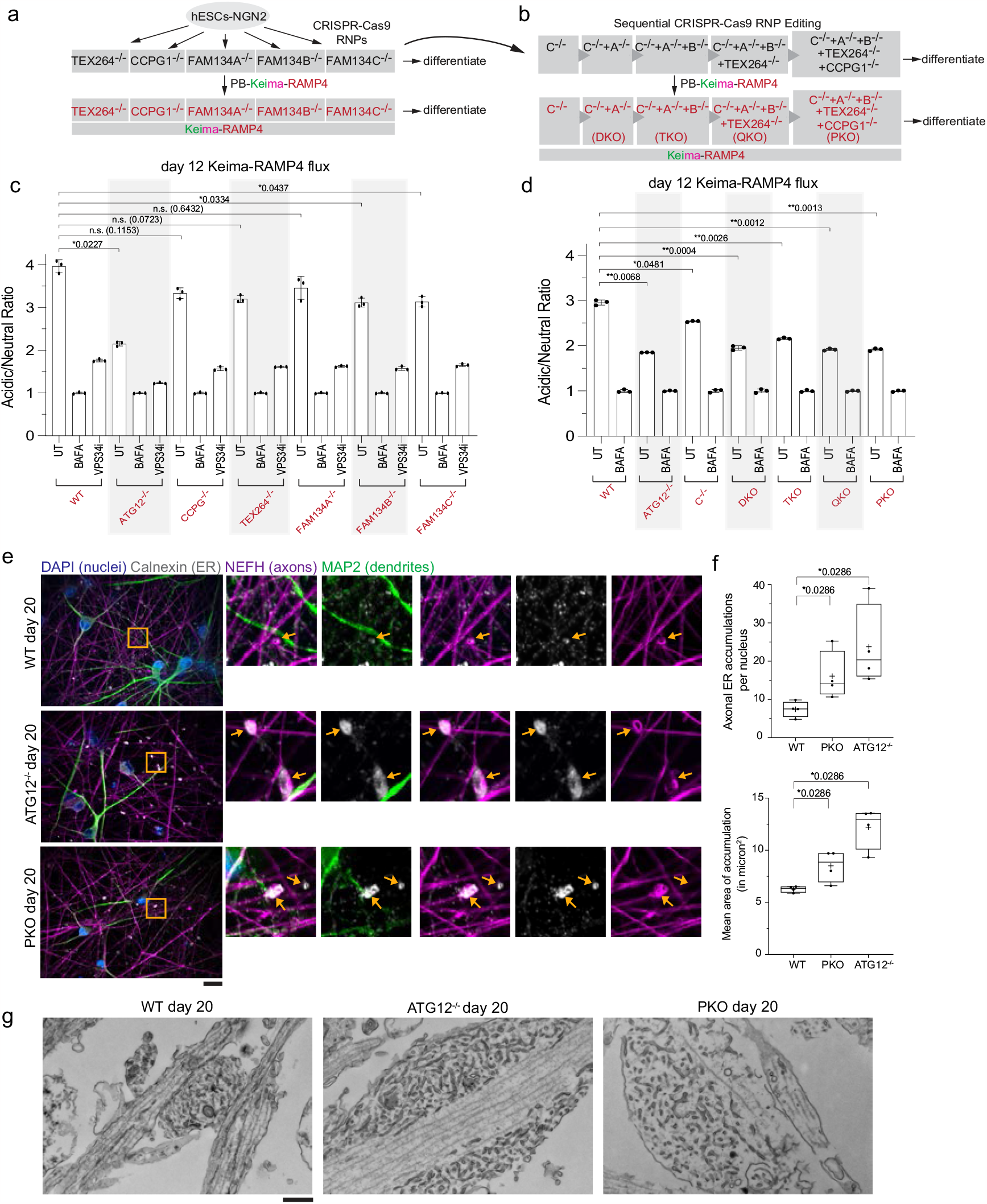
Combinatorial regulation of ER clearance via ER-phagy receptors during neurogenesis in vitro. **a-b**, A toolkit for analysis of ER-phagy receptors. hESCs were subjected to CRISPR-Cas9 gene editing to delete individual (panel **a**) or multiple (panel **b**) receptors. Keima-RAMP4 was expressed in each of the mutant hESCs, prior to analysis during differentiation. **c-d**, Ratiometric analysis of Keima-RAMP4 flux in the indicated WT or mutant hESCs was measured by flow cytometry at day 12 of differentiation. The ratio of acidic to neutral Keima fluorescence was normalized to samples treated with BAFA (100 nM) for 4 h. Each measurement reflects biological triplicate measurements. Error bars represent SD. *, p<0.05; **, p<0.01; n.s., not significant; Brown-Forsythe and Welch One-way ANOVA and Dunnett’s T3 multiple comparisons test. **e-f**, PKO iNeurons accumulate aberrant ER structures, particularly in axons. WT or day 20 iNeurons of the indicated genotypes were immunostained with α-Calnexin (ER, white), α-MAP2 (dendrites, green), α-NEFH (axons, magenta), and with DAPI (nuclei, blue). Scale bar, 25 microns. Number of axonal ER accumulations/nucleus (panel **f, top**) or mean area of ER accumulation (panel **f, bottom**) is represented with min-to-max box-and-whiskers plots. Lines are at medians and “+” symbols designate the means. Four points shown for each WT or KO condition represent the measured values from four independent differentiations. *, p<0.05; Mann-Whitney test. **g**, TEM images of sections though WT, ATG12^-/-^ and PKO axons containing enlarged structures containing areas of ER membranes. Scale bar, 500nm.

### ER-phagy receptor control of ER-phagic flux in iNeurons

To directly examine individual receptor contribution to ER-phagy during differentiation, we measured Keima-RAMP4 ER clearance to the lysosome in receptor mutant cells at day 0, 4, or 12 of differentiation using flow cytometry (**Fig. 5c, Extended Data Fig. 6f,g**). As expected, Keima-RAMP4 flux increased from 2.5 to 4.0-fold in WT cells at day 4 and 12 of differentiation, which was substantially reduced in day 12 ATG12^-/-^ iNeurons (**Fig. 5c, Extended Data Fig. 6f,g**). In contrast, all single mutants displayed Keima-RAMP4 flux comparable to WT at day 4 and >80% of WT at day 12 (**Fig. 5c, Extended data Fig 6g**). However, upon elimination of FAM134A/C, the level of Keima-RAMP4 flux was reduced from WT, approaching a similar level to that seen with ATG12^-/-^ cells at day 12 of differentiation, with a slight further reduction upon removal of additional receptors in the context of this pan-ER reporter (**Fig. 5d**). In these flow cytometry experiments, we gated for live single cells. Each genetic background had a similar percent of live cells, which emphasizes that the reduction in ER phagic flux measured for ATG12^-/-^ and PKO iNeurons was not skewed due to differential cell viability (**Extended Data Fig. 6h, source data contains flow cytometry gating strategy**).

Consistent with defective ER turnover, day 20 PKO iNeurons displayed more abnormally enlarged α - Calnexin marked ER structures in α -NEFH-positive axons (**Fig. 5e**). The number and size of these structures were intermediate between WT and ATG12^-/-^ iNeurons, as visualized by light microscopy (**Fig. 5f**). TEM of thin sections through axons of PKO iNeurons also revealed examples of frequent dilated structures rich in tubular ER, similar albeit smaller than that observed in ATG12^-/-^ axons (**Fig. 5g; Extended Data Fig. 6i**).

### Combinatorial receptor control of ER-proteome remodeling in iNeurons

While results thus far revealed a prominent role for FAM134A/C in pan-ER Keima reporter flux during neurogenesis, the identities of proteins subject to remodeling during differentiation and the specificity of ER clearance via individual receptors were unclear. Although many individual ER proteins display small 1.2 to 1.5-fold (Log_2_FC from 0.26 to 0.58) increases in abundance in ATG12^-/-^ iNeurons, the ER compartment accumulates when viewed as a cohort (**Fig. 1c**). We were further curious as to the distribution of these fold changes and wondered why some ER proteins accumulated more than others. This gradient in accumulation opened up the possibility that certain ER proteins may be enriched effectively in regions of the ER that are undergoing autophagic capture by ER-phagy receptors, reflecting either sub-organelle localization or a propensity to exist at highly curved membranes associated with ER-phagic capture^5,6,20-22^. To unmask any potential selectivity of ER-phagy receptors for specific clients, we performed 18-plex TMT quantitative proteomics using single (**Fig. 6a, Supplementary Table 5)** and combinatorial (**Fig. 6b, Supplementary Table 3**) ER-phagy receptor mutants at day 12 of differentiation. As a comparator, ATG12^-/-^ iNeurons were included as a control for autophagy-dependent stabilization. Abundance of organelles at the global level, including ER, was largely unaffected in single ER-phagy mutants, as suggested by violin plots for individual organelle proteomes (**Fig. 6a, Extended Data Fig. 7a**). In contrast and consistent with a more pronounced effect on Keima-RAMP4 flux and axonal ER accumulation, combinatorial mutants displayed an overall increase in ER protein abundance comparable to that seen in ATG12^-/-^ (**Fig. 6b-c, Extended Data Fig. 7b-d)**. The distribution of ER proteins in ATG12^-/-^ or the DKO to PKO mutants significantly deviate from a randomized selection of proteins (randomized control) of the same number of proteins (**Fig. 6c**). However, the combinatorial mutants did not affect the distribution of the Golgi proteome know to be regulated by loss of ATG12 in this system^26,38^, which is consistent with a specific role for these ER-phagy receptors in ER turnover **(Extended Data Fig. 7b-d**). Importantly, we confirmed using quantitative proteomics that ER protein accumulation was maintained in later stage day 20 ATG12^-/-^ and PKO iNeurons corresponding to the time employed for many of the imaging experiments described above, and we confirmed this ER accumulation occurs in an independent PKO clone (**Extended Data Fig. 8a-c; Supplementary Table 6**).

**Fig. 6.**
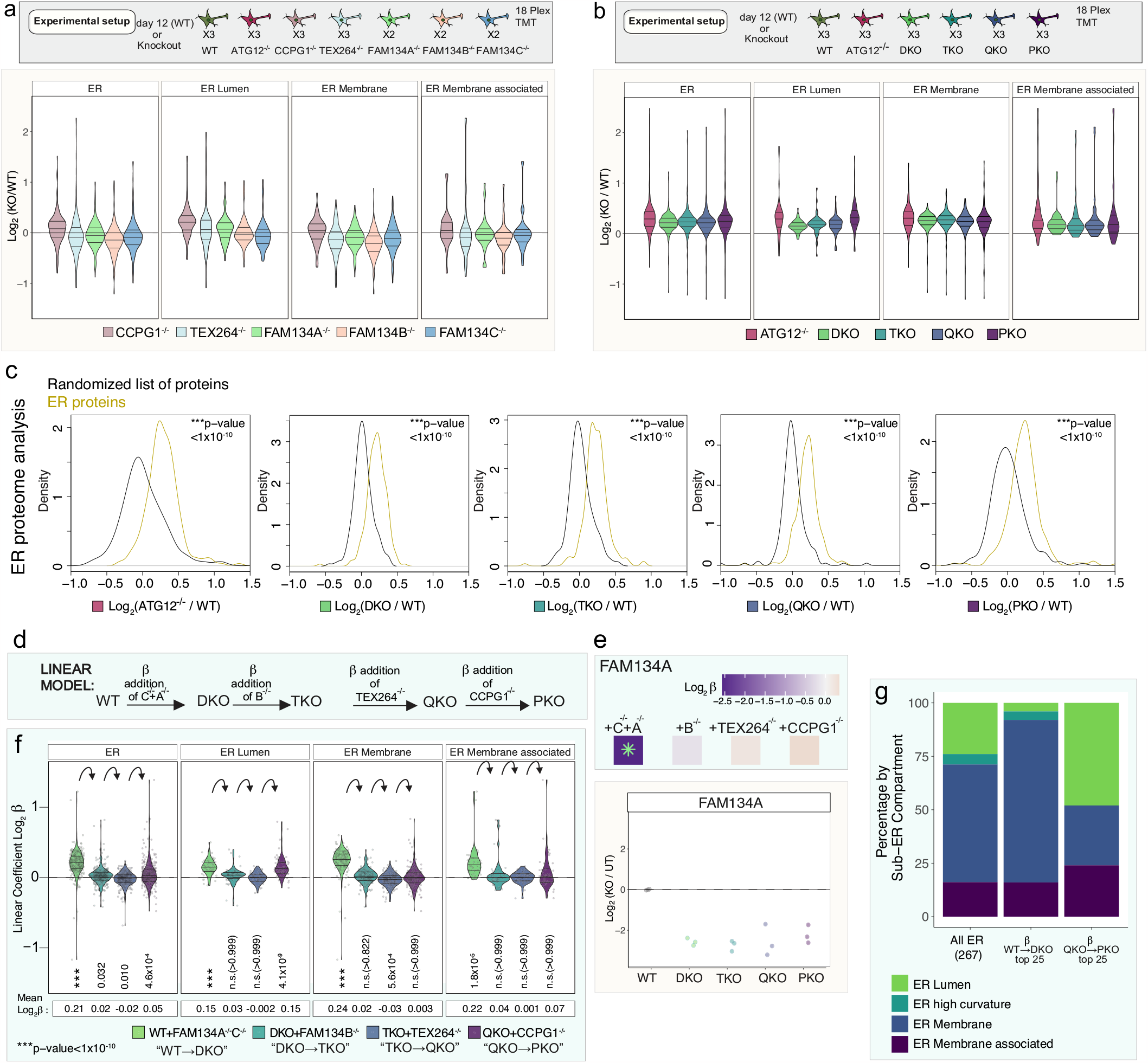
Selectivity of ER-phagy receptors in ER remodeling in iNeurons revealed by combinatorial multiplexed proteomics. **a**, Scheme depicting an 18-plex TMT experiment examining total proteomes of the indicated single ER-phagy receptor mutant day 12 iNeurons. Violin plots (lower panel) depicting Log_2_FC (mutant/WT) for the indicated classes of ER proteins in single mutant iNeurons (day 12) are shown in the lower plot. **b**, Scheme depicting an 18-plex TMT experiment examining total proteomes of the indicated combinatorial ER-phagy receptor mutant day 12 iNeurons. Violin plots (lower panel) depicting Log_2_FC (mutant/WT) for the indicated classes of ER proteins in combinatorial mutant iNeurons (day 12) are shown in the lower plot. **c**, Log_2_FC (mutant/WT) distributions of ER proteins compared to randomized selections of the same number of proteins (100 iterations). p-values for each comparison are calculated with a Kolmogorov-Smirnov Test (two-sided). **d**, Application of a linear model to identify selective cargo for individual ER-phagy receptors via quantitative proteomics. In the linear model, a coefficient FC (β) is calculated for sequential loss of ER-phagy receptors starting from WT to DKO, then DKO to TKO, then TKO to QKO, then QKO to PKO. **e**, Top panels (β coefficient values) and lower panels (Log_2_FC) for FAM134A. Green asterisk in top panel indicated significant change (adjusted p-value <0.05) in β coefficient for that mutant. This analysis is distinct from traditional comparisons between each mutant and WT (lower panel). **f**, Violin plots depicting β coefficient FC for the indicated classes of ER proteins. p-values for each comparison are calculated with a Wilcoxon test with Bonferroni correction. **g**, Top 25 accumulating ER proteins in WT to DKO and QKO to PKO and their respective ER compartment compared to the landscape of the whole ER.

In cancer cell lines, MTOR inhibitor Torin1 induces a starvation-like response, leading to clearance of ER (among other organelles) and proteins via autophagy, but this clearance is blocked in autophagy deficient cells^39^.To further probe susceptibility of ER to selective turnover via general autophagy as compared with selective ER-phagy, we examined organelle and proteome abundance in iNeurons treated with Torin1 for 15 h. In ATG12^-/-^ iNeurons, organelle clearance was blunted when compared with WT iNeurons, which is consistent with autophagic turnover being blocked (**Extended Data Fig. 8d, Supplementary Table 7**). Consistent with findings that the PKO primarily affects only ER proteome remodeling, PKO iNeurons treated with Torin1 demonstrated a defect in clearance of ER proteins (similar to ATG12^-/-^), while other organelles were largely unaffected (in contrast to ATG12^-/-^) (**Extended Data Fig. 8d, Supplementary Table 7**). The ubiquitin-binding autophagy receptor CALCOCO1 has been reported to function as a soluble receptor for turnover of both Golgi and ER in response to nutrient stress ^40,41^, although a general function in ER- or Golgiphagy has been brought into question ^38^. While CALCOCO1 accumulated in ATG12^-/-^ iNeurons during differentiation (**Fig. 1c**), we found that iNeurons lacking CALCOCO1 displayed no global accumulation of ER or Golgi proteomes, unlike ATG12^-/-^ iNeurons examined in parallel by TMT proteomics (**Extended Data Fig. 8e-i, Supplementary Table 8**). Thus, CALCOCO1 alone is not necessary for ER or Golgi maintenance in this system.

### Quantitative modeling of ER proteome remodeling via ER-phagy

The behavior of the ER proteome in single and combinatorial ER-phagy mutant iNeurons (**Fig. 6a-b, Extended Data Fig. 9a**,**b**) suggests both redundancy and selectivity for client turnover by receptors, with a range of alterations in abundance observed across the ER proteome (**Fig. 6a-b, Extended Data Fig. 9a,b**). To further probe this underlying specificity, we employed a linear model that measures the sequential effect of (1) FAM134A/C, (2) FAM134B, (3) TEX264, and (4) CCPG1 deletion. The model measures the positive or negative Log_2_FC values comparing each step and assigns these changes with positive or negative β coefficients [(1) β^WT→DKO^; (2) β^DKO→TKO^, (3) β^TKO→QKO^, and (4) β^QKO→PKO^, (**Fig. 6d, Extended Data Fig. 10a,b**) (see **METHODS**). The behavior of FAM134A is exemplary of the applicability of the model (**Fig. 6e**): β^WT→DKO^ was strongly negative (−2.5) and significant (indicated by the asterisk), consistent with its deletion in the DKO mutant compared to WT, but β coefficient values in subsequent deletions was near zero and not significant, as expected since FAM134A remains deleted and therefore remains at the same abundance throughout the remainder of the allelic series.

Global analysis revealed an increase in mean β^WT→DKO^ coefficients for the ER proteome (0.21), which was primarily reflected in alterations in the abundance of ER-membrane and ER-lumen proteins (**Fig. 6f**). β^DKO→TKO^ and β^TKO→QKO^ coefficients reflecting the further deletion of FAM134B and TEX264, respectively, are near zero for the ER proteome as a whole and for specific ER subregions (**Fig. 6f, Extended Data Fig. 10a**), suggesting modest or no contributions to ER turnover in this context. In contrast, the mean β^QKO→PKO^ coefficient resulted in an increase (0.15) for a cohort of ER lumenal proteins (**Fig. 6f, Extended Data Fig. 10a**), indicating that CCPG1 and FAM134A/C independently control the abundance of a set of lumenal proteins based on either the magnitude of abundance change or protein identity, as explored further below. The effect of CCPG1 on lumenal ER protein abundance is further demonstrated by organelle point plots comparing β^TKO→QKO^ and β^QKO→PKO^, with significant displacement of ER lumen off the diagonal (**Extended Data Fig. 10a**). We next compared how each organelle proteome changes when an individual deletion is made in wildtype iNeurons to the organelle proteome changes that occur when the same deletion is added to the sensitized background reflected by the β value for that deletion in the combinatorial deletion series (**Extended Data Fig. 10c**) The effect on the ER by single deletion of CCPG1 or by adding CCPG1 to the QKO to create the PKO suggests CCPG1 can act alone as an ER-phagy receptor to clear luminal proteins during neuronal differentiation. However, the FAM134 family of receptors only yielded an increase in the ER network and different ER compartments when the FAM134 family was deleted in combination.

The finding that combined loss of FAM134A and C leads to accumulation of a cohort of ER proteins and that the ER proteome was not substantially altered upon further deletion of FAM134B led us to ask whether FAM134A and B were functionally equivalent in this setting. We generated FAM134B/C^-/-^ cells and performed multiplexed proteomics comparing FAM134C^-/-^, FAM134A/C^-/-^ (DKO), and FAM134B/C^-/-^ iNeurons (day 12) (**Extended Data Fig. 10d, Supplementary Table 9**). Global ER and particularly ER-membrane protein abundance also increased in FAM134B/C^-/-^ iNeurons relative to FAM134C^-/-^ iNeurons (**Extended Data Fig. 10d**). Taken together, this suggests that FAM134 copy number, rather than the identity of the specific isoform, underlies ER proteome remodeling in this context.

### ER-phagy receptor substrate specificity

To directly examine substrate selectivity of ER-phagy receptors, we first explored the top 25 ranked proteins with positive β coefficients for both β^WT→DKO^ and β^QKO→PKO^. When compared with all ER proteins, those with positive β^WT→DKO^ coefficients were particularly enriched in ER membrane proteins while in contrast, proteins with positive β^QKO→PKO^ coefficients were enriched in lumenal proteins (**Fig 6g**). Heatmaps revealing the identity of these top accumulators highlight the degree of change in β coefficients, with significantly changing proteins marked with an asterisk (*adjusted p-value <0.05; positive or negative β coefficients, **Fig. 7a**). The extent of accumulation of these top ranked proteins in PKO cells was similar to that seen with ATG12^-/-^ iNeurons (**Fig. 7a**), indicating that the PKO mutant closely approximates the biochemical phenotype of ATG12 deficiency for ER turnover. Globally, we identified 84 membrane proteins with significantly (*adjusted p-value <0.05) positive or negative β ^WT→DKO^ coefficients, which were distributed across multiple functional categories and contained varying numbers of TM segments (**Fig. 7b and Supplementary Table 3**). Given that several of the ER shaping proteins with RHDs are within this group of significant changers (**Fig. 7b**) and ATG12 deficiency strongly affects ER shaping proteins with RHDs (**Fig. 1e**), we examined this class of proteins further. In addition, we examined ER proteins that are specifically related to two neurological disorders, hereditary spastic paraplegia (HSP) and hereditary sensory and autonomic neuropathy (HSAN) (a subset of which are also ER curvature shaping proteins that contain RHDs). Heatmaps of Log_2_FCs values for these specific ER proteins are found in **Extended Data Fig.11a,b**), and immunoblotting of selected proteins confirmed accumulation both in ATG12^-/-^ and PKO iNeurons (**Extended Data Fig. 11c)**. First, we found that a subset of ER curvature proteins specifically increased in β ^WT→DKO^ including RTN1-C (Log_2_FC=0.44) (**Fig. 7a-c, Extended Data Fig.11c,d)**. Similarly, REEP5 accumulated – albeit to a lesser extent – in PKO iNeurons (**Fig. 7b-c, Extended Data Fig.11a,c**), and Keima-REEP5 flux measurements revealed decreased flux in PKO cells, approaching that observed in ATG12^-/-^ iNeurons (**Fig. 7d)**. Second, a distinct set of RHD proteins (REEP1, REEP3, REEP4) decrease in abundance, and display negative β coefficients for DKO (**Fig. 7a,b,e** and Extended Data Fig. 11a-c,e). REEP1 also further decreases upon deletion of TEX264, as indicated by a significant negative β -coefficient and Log_2_FC (**Fig. 7e**). Since members of the RHD protein family (e.g. REEP1) are strongly upregulated during iNeuron differentiation (**Fig. 1a-d, Extended Data Fig. 11f**), alterations in abundance across the REEP family indicate distinct pathways for controlling ER shape remodelling for neurons specifically via ER-phagy. Whereas the collective ER proteome did not increase for the single FAM134C deletion, abundance alterations for ER-shaping proteins specifically were observed with just the single deletion (**Extended Data Fig. 11a**), indicating that FAM134C likely contributes substantially to differential regulation of shaping proteins during neurogenesis (**Fig. 7c**). Interestingly, ATG12^-/-^ iNeurons display increases in abundance for all REEP proteins, indicating that a broad block to autophagy can mask otherwise distinct proteome remodeling events relevant to an individual ER-phagy receptor (**Extended Data Fig. 11a-c,f**).

**Fig. 7.**
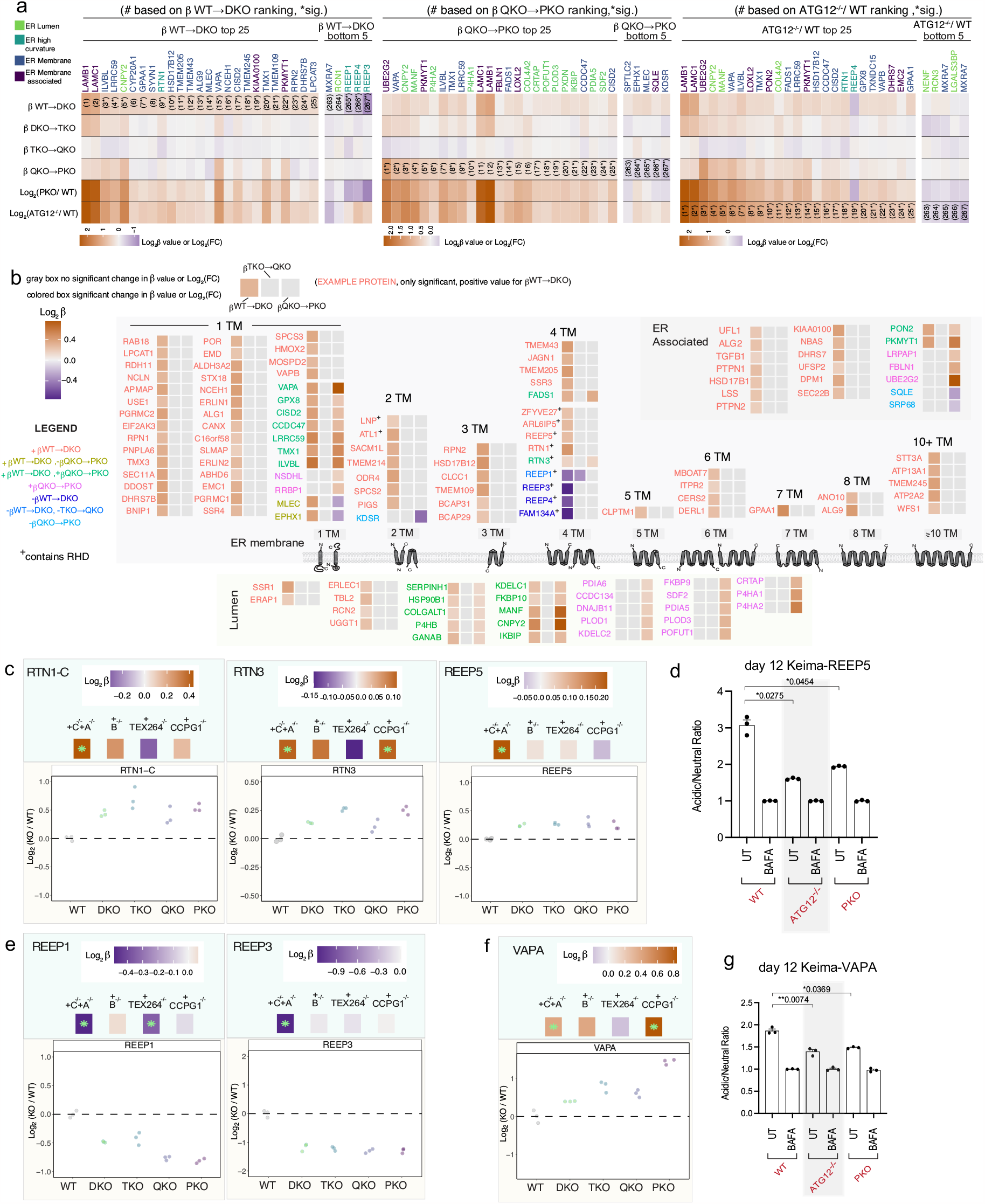
ER-phagy receptor remodeling of the ER proteome landscape and ER-phagy receptor cargo specificity during iNeuron differentiation. **a**, Top 25 accumulated and bottom 5 depleted ER proteins ranked on WT to DKO β coefficient values (left panel), QKO to PKO β coefficient values (middle panel), or on Log_2_FC (ATG12^-/-^/WT). **b**, The ER-associated, ER membrane, or ER lumenal distribution and predicted trans membrane character of ER proteins with significant β coefficient values (*, adjusted p-value <0.05) in WT to DKO (111 up, 4 down), TKO to QKO (1 down) and QKO to PKO (39 up, 5 down). Zero proteins were significant in DKO to TKO. Each protein name is colored based on if there is a significant change in these steps in the allelic series as according to the legend. The corresponding β coefficient value heatmap for each protein is colored in if there is a significant change and left blank if there is no significant change at that step in the allelic series (see legend). **c**, Examples of ER shaping proteins with significant β coefficients that accumulate at one or more steps in the allelic series. Top panels (β coefficient values) and lower panels (Log_2_FC) for single proteins including RTN1-C, RTN3, REEP5. **d**, Keima-REEP5 flux measurements in WT, ATG12-/-, and PKO iNeurons (day 12) using acidic/neutral ratios in the presence of BAFA for normalization. **e**, As in **c**, but for proteins REEP1 and REEP3 (ER shaping proteins with significant β coefficients that decrease). **f**, As in **c** and **e** but for VAPA (ER membrane protein that forms contact sites with other organelles). **g**, Autophagic flux assay for Keima-VAPA in WT, ATG12^-/-^ or PKO iNeurons (day 12). For individual protein plots in **c, e**, and **f** the green asterisks in the top panels indicate significant change (*, adjusted p-value <0.05) in β coefficients for each mutant. For autophagic flux experiments using Keima-REEP5 and Keima-VAPA, n = 3. *, p<0.05; **, p<0.01; Brown-Forsythe and Welch One-way ANOVA and Dunnett’s T3 multiple comparisons test.

To examine whether loss of specific REEP family members is directly due to loss of FAM134, we ectopically expressed FAM134C-GFP in either WT or TKO ES cells using a PiggyBac vector and converted the cells to iNeurons (day 12) (**Extended Data Fig. 11g-i**). Immunoblotting of cell extracts revealed that the increase in TEX264 abundance and the decrease in REEP1 or REEP4 in TKO cells is reversed by reintroduction of FAM134C-GFP (**Extended Data Fig. 11g,i**). Similarly, proteomics revealed that FAM134C-GFP expression in TKO iNeurons reversed the global accumulation of a cohort of ER proteins, in particular ER-membrane proteins (**Extended Data Fig. 11h, Supplementary Table 10**). Proteomics also validated the rescue in expression of REEP1/2/3; REEP4 was not detected in this specific experiment (**Extended Data Fig. 11i**).

The ER lumenal compartment is primarily responsible for folding and modification of secretory and membrane proteins, but proteins in this compartment have also been reported to undergo autophagic trafficking^9,42,43^. We identified two major patterns of ER lumenal protein abundance changes, reflected in β^WT→DKO^ and β^QKO→PKO^ coefficients. In total, 25 ER lumen proteins (primarily lacking a TM) were stabilized in the DKO mutant, and 10 these were further stabilized in PKO mutants (**Fig. 7b and Supplementary Table 3**). In contrast, a distinct cohort of 16 ER lumen proteins was stabilized specifically in the PKO mutant with no significant effect observed with DKO, TKO, or QKO mutants (e.g. P4HA1 and P4HA2) (**Fig. 7a,b, Extended Data Fig. 12a,b and Supplementary Table 3**), and Log_2_FC for these lumenal proteins were also stabilized in ATG12^-/-^ iNeurons (**Extended Data Fig. 12b**). These findings suggest both redundant and specific lumenal cargo for FAM134A/C and CCPG1 receptors. We also found it compelling that deletion of CCPG1 alone or in the context of a TEX264^-/-^/CCPG1^-/-^ double mutant resulted in increased abundance of a subset of lumenal proteins with significant similarities to that seen with the PKO mutant (**Extended Data Fig. 10c, 12b,c and Supplementary Table 11**).

Intriguingly, the single TM segment proteins VAPA and VAPB, which mediate contact site interactions between ER and a number of other organelles, including mitochondria, via an interaction with VPS13 and other lipid transfer proteins^44^, have a positive β coefficient in DKO and/or PKO mutants, indicating that VAPs undergo multiple modes of ER-phagic turnover (**Fig. 7a,b,f and Extended Data Fig. 12d**). An increase in VAPA abundance was also observed by immunoblotting in both ATG12^-/-^ and PKO iNeurons (**Extended Data Fig. 11c**). To directly examine the hypothesis that VAPA is an ER-phagy client, we created a Keima-VAPA reporter construct which was expressed via PiggyBac in WT, ATG12^-/-^ and PKO iNeurons. We found that Keima-VAPA autophagic flux was reduced in PKO iNeurons to an extent similar to that seen with ATG12^-/-^ iNeurons, consistent with the idea that VAPA is a substrate for ER-phagy (**Fig. 7g**). VAPA abundance was also increased ATG5^-/-^ cerebellar granule neurons in culture^23^. In parallel, β coefficient correlation plots for organelles reveals selective accumulation of mitochondria as a result of CCPG1 deletion (**Extended Data Fig. 7c, 10a-c)**. These findings open further study into how ER-phagy mechanisms are regulating ER architecture to facilitate functions like maintaining robust yet dynamic contact sites with other organelles.

## .DISCUSSION

Previous studies indicated that loss of autophagy pathways in mice or ES cells leads to increased accumulation of ER proteins^23,26^, but the extent to which this reflects non-specific macroautophagy or selective ER-phagy was unknown. Indeed, RNAi mediated suppression of individual ER-phagy receptors failed to promote axonal ER accumulation in cultured mouse neurons, leading to the suggestion that ER-phagy may be a constitutive process linked with autophagosome formation in distal axons^23^. The use of a genetically tractable iNeuron system, which displays a dramatic accumulation of axonal ER in the absence of a functional autophagy system^26^, has allowed us to examine roles for multiple ER-phagy receptors during ER remodeling associated with neurogenesis.

Using live-cell imaging, we found that these FAM134C and TEX264 are mobilized into LC3B-positive vesicles that traffic in axons. Previous studies in the context of nutrient stress demonstrated that FAM134 and TEX264 are concentrated into the same ER structures that are captured during ER-phagy while CCPG1 also undergoes ER-phagy but forms distinct ER-cargo domains^10^. Current models indicate that ER serves as a source of lipids for phagophore formation but that ER membrane proteins themselves are not incorporated into autophagosomal membranes^24,25^. Thus, we conclude that FAM134C and TEX264-positive puncta reflect ER-phagy rather than the process of autophagosome biogenesis as previously observed in distal axons^36^. Consistent with this, we readily observed LC3-positive autophagosomes that traffic on microtubules but lack detectable TEX264 or FAM134C proteins, indicative of distinct cargo (**Fig. 3e**). One plausible cargo is mitochondria, which is selectively incorporated into a subset of autophagosomes in mouse neurons^45^. Importantly, we observed numerous autophagosomes in axons via cryo-ET, some of which contain ER membranes and TEX264-GFP in correlative imaging. Tomogram reconstruction revealed the presence of membranes consistent with tubular ER (**Fig. 4**).

Generation of single and combinatorial ER-phagy receptor knockouts complimented with general ER flux measurements and quantitative proteomics revealed that selective ER-phagy mechanisms clear ER proteins during iNeuron differentiation. Mutation of FAM134A and C or FAM134 B and C was sufficient to produce a global increase in the ER proteome, with the transmembrane ER proteome featured prominently among the most stabilized proteins. In contrast, deleting CCPG1 in different allelic backgrounds revealed CCPG1’s primary role in clearing lumenal proteins. Unlike FAM134 family members and TEX264, CCPG1 contains a lumenal domain that has been suggested to associate with lumenal autophagy substrates^9,42,46^. Our proteomic analysis validates previously reported CCPG1 cargo (e.g. P3H4)^42^ and provides additional candidates for further analysis. Unlike ER-phagy in response to nutrient stress ^10,11^, it does not appear that loss of TEX264 alone affects ER network clearance during neurogenesis and our results suggest that TEX264 ER-phagic clearance is dependent on the FAM134 ER-phagy receptor family in this context.

The RHD domains of FAM134 family members have been proposed to cluster into highly curved membranes during an early step in ER-phagy initiation, thereby promoting ER membrane budding and scission of ER membrane into autophagosomes for subsequent lysosome degradation.^16,22^ Other RHD-containing proteins, including REEP5 and ARL6P1/5,^22,47^ can associate with FAM134C in co-immunoprecipitation experiments. We found distinct behavior in the abundance of various RHD proteins. While the majority of REEP proteins and several RTN proteins accumulated in ATG12^-/-^ iNeurons, cells lacking FAM134A/C accumulate REEP5 and RTN1 but display loss of REEP1-4. REEP1-4 protein abundance was rescued upon expression of FAM134C-GFP in FAM134A/C^-/-^ cells. One possible explanation for this result is that FAM134 proteins facilitate REEP1-4 trafficking or stability. Unlike REEP5/6, which contain 4 reticulon helices within the outer leaflet of the ER membrane, REEP1-4 contain only 3 such helices and therefore may have distinct functional properties. In yeast, the REEP1-4 ortholog YOP1 has been shown to bind to highly curved membranes, including small vesicles, in contrast to REEP5/6.^47^ Future studies are required to understand the distinct properties of REEP proteins observed here. Upstream signals such as phosphorylation are known to activate and possibly cluster multiple ER-phagy receptors^9,48,49^, but further studies are required to understand the relevant regulatory pathways important for axonal ER-phagy and membrane remodeling associated with autophagosomal capture.

ER tubule proteins shape the ER network and facilitate dynamic connections between ER and other organelles, which links each organelle compartment of this highly polarized neuronal cell system to maintain optimal function. Importantly, several proteins found to be controlled via selective autophagy are either linked with human disease (e.g. mutation in ER contact site protein VAPB results in motor neuron disease), or, when experimentally deleted, result in altered ER structure within neurons with functional consequences^50^. This work provides a versatile resource for further interrogating how ER remodeling is optimized for various cell states via selective ER-phagy.

## ACKNOWLEDGMENTS

This work was supported by a Harvard Medical School sponsored research project grant from Astellas Pharmaceuticals (to J.W.H.), from Aligning Science Across Parkinson’s (ASAP, Michael J Fox Foundation administers the grant ASAP-000282 on behalf of ASAP and itself to J.W.H., B.A.S., W.B.), NIH (R01NS083524, R01NS110395 to J.W.H., R01GM132129 to J.A.P.), the Max Planck Society (B.A.S.), and the Harvard Medical School Cell Biology Initiative for Molecular Trafficking and Neurodegeneration. M.J.H. was supported by post-doctoral fellowships from Jane Coffin Childs and the Fred and Joan Goldberg Education Fund at Harvard Medical School. Michael J Fox Foundation administers the grant ASAP-000282 on behalf of ASAP and itself. For the purpose of open access, the author has applied a CC-BY public copyright license to the Author Accepted Manuscript (AAM) version arising from this submission. We also thank the Harvard Electron Microscopy Core and the Nikon Imaging Center at Harvard Medical School for microscopy support.

## AUTHOR CONTRIBUTIONS

M.J.H. and J.W.H. conceived the study. M.J.H. performed proteomics, gene editing, cell line construction, biochemical assays, cell biology, imaging, and informatic analyses. J.C.P. and Y.J. performed gene editing and cell line construction. I.R.S. and M.G.-L. performed informatic analysis of proteomic data. J.A.P. provided proteomics support. C.C. and A.B. performed cryo-ET under the supervision of F.W., W.B., and B.A.S. The manuscript was written by J.W.H., M.J.H., C.C. and I.R.S. with input from all authors.

---

**Extended Data Fig. 1.**
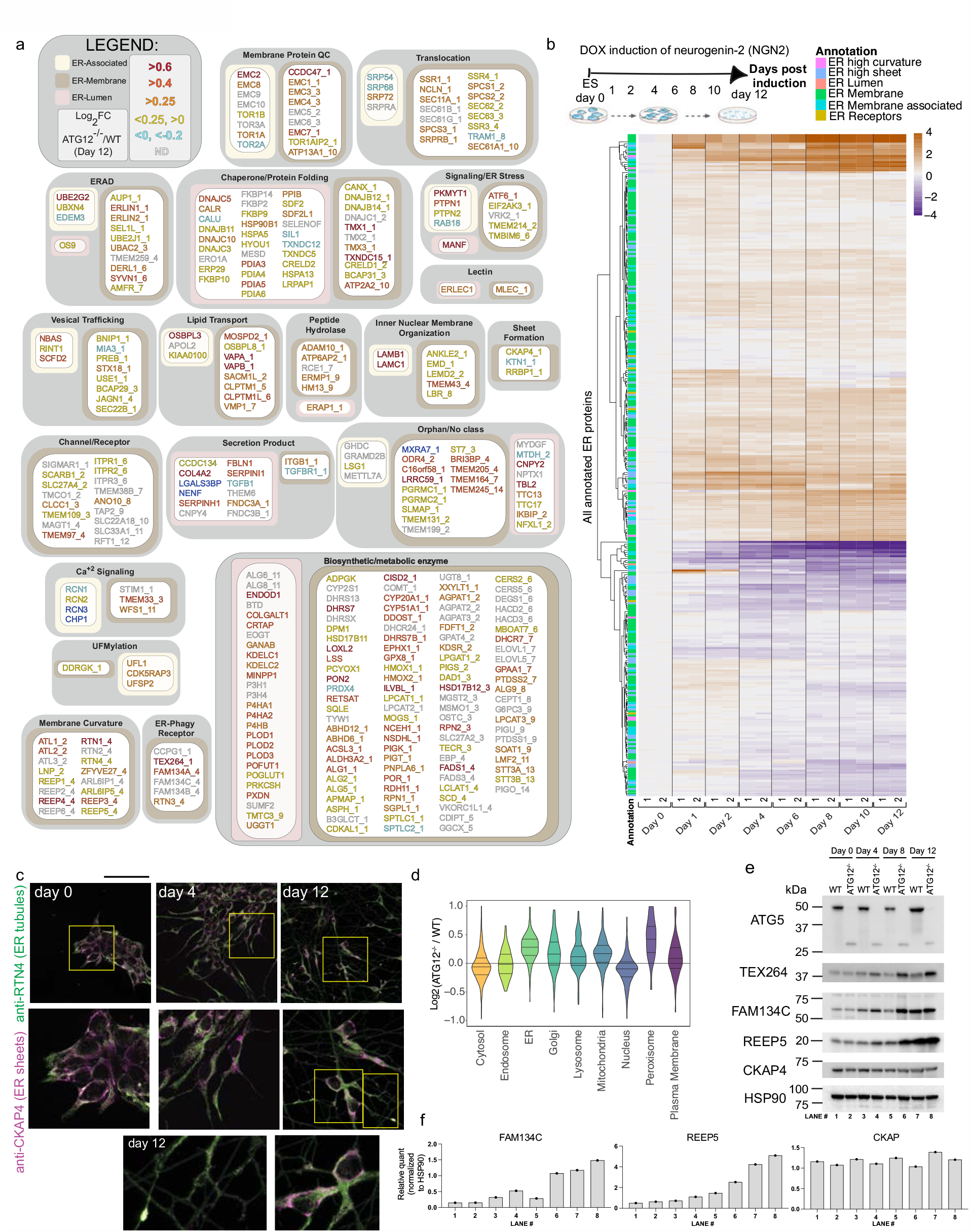
Landscape of ER remodeling via autophagy during hESC differentiation to iNeurons in vitro. **a**, Landscape of the ER proteome and the effect of autophagy on accumulation of individual proteins. The ER proteome (359 proteins, **Supplementary Data Table 1**) is organized into functional modules and protein attributes (involved in ER membrane curvature, ER-associated, ER-membrane, ER-Lumen or ER-phagy receptor) are indicated by the respective outline box color (see inset legend). For proteins with transmembrane segments, the number of segments are indicated after the protein name (_1, _2, etc) based on data in Uniprot. The text of each protein name is colored based on day12 ATG12^-/-^ vs WT Log_2_FC (see inset legend). (**Supplementary Data Table 3**). **b**, Changes in the abundance of the ER proteome (267 detected proteins) during conversion of WT hESCs to iNeurons are shown in as heatmaps (Log_2_FC) at the indicated day of differentiation relative to hESCs. Data are from our previous analysis of iNeuron differentiation. Annotations of the type of ER protein are indicated by the relevant colors. **c**, hESCs were differentiated to iNeurons and stained with antibodies against CKAP4 enriched in ER sheets (magenta) and RTN4 enriched in ER-tubules (green) at day 0, 4 and 12 of differentiation. RTN4 staining is evident throughout neuronal projections. Scale bar, 100 microns. **d**, Violin plots for relative abundance of proteins located in the indicated organelles in ATG12^-/-^ versus WT day 12 iNeurons. **e**, Immunoblots of cell extracts from WT or ATG12 ^-/-^ hESCs for the indicated day of differentiation. Blots were probed with the indicated antibodies, with α-HSP90 employed as a loading control.

**Extended Data Fig. 2.**
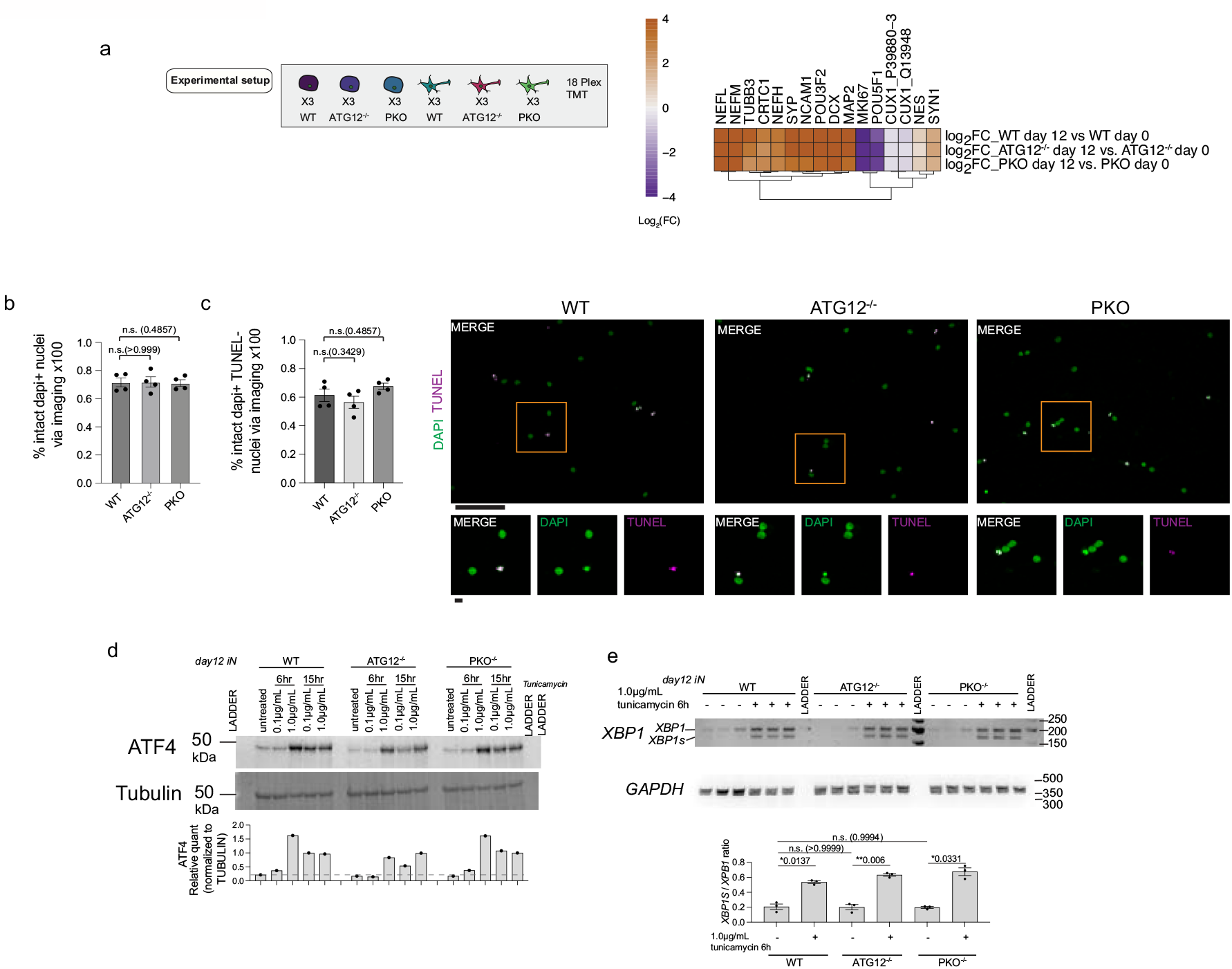
Quality control of in vitro neurogenesis methods. **a**, The indicated ES cells were either cultured in the pluripotent state (day 0) or converted to day 12 iNeurons prior to total proteome analysis by multiplex TMT in biological triplicate cultures (n=3). The relative abundance (Log_2_ FC) of the indicated neurogenesis or pluripotency factors at day 12 relative to day 0 is shown, indicating that all genotypes undergo differentiation to a similar extent as measured by markers of the process. **b**, Comparable number of viable iNeurons were observed upon quantitative analysis of intact DAPI-positive nuclei compared to all DAPI-positive (intact and fragmented) DNA structures in cultures of the indicated genotypes. Four points shown for each WT or KO condition represent the measured ratios from the same four independent differentiations also analyzed for ER structure per nuclei in **Fig 2b**. *, p<0.05, Mann-Whitney test. **c**, Quantification of DAPI -positive TUNEL-negative nuclei in day 20 iNeurons from the indicated genotypes and representative images of DAPI-positive nuclei in green and TUNEL-stained DAPI-positive fragmented nuclei, with TUNEL signal in magenta. The lower panels are magnified regions (merges and separate image channels) of the area boxed in the respective image above. Four points shown for each WT or KO condition represent the measured values from four independent differentiations. *, p<0.05, Mann-Whitney test. Scale bar, 100 microns (top), 10 microns (bottom) **e**, ATF4 abundance was examined by immunoblotting of extracts from day 12 iNeurons from the indicated genotypes with or without treatment with tunicamycin as a positive control for induction of the ER-stress stress response. Tunicamycin was used at 0.1 or 1.0 μg/ml for 6 or 12 h, as indicated. Blots were reprobed with α-tubulin as a loading control. The relative α-ATF4 signal, normalized for tubulin, is shown in the histogram (lower panel) with the dashed line representing ATF4 signal in untreated WT cells. n=1. **f**, *XBP1* and *XBP1s* mRNA from the indicated iNeurons in biological triplicate (n=3) cultures was subjected to reverse transcription-PCR (see **METHODS**) and examined by agarose gel electrophoresis to resolve spliced and unspliced *XBP1*. 1.0 μg/ml tunicamycin (6h) was employed as a positive control for induction of ER stress and *XBP1* splicing. *GAPDH* was used as a positive control. The *XBP1s*/*XBP1* ratio was quantified as shown in the histogram (lower panel). No evidence of increased *XBP1* splicing was observed in any genotype under untreated conditions. *, p<0.05; **, p<0.01; n.s., not significant; Brown-Forsythe and Welch One-way ANOVA and Dunnett’s T3 multiple comparisons test.

**Extended Data Fig. 3.**
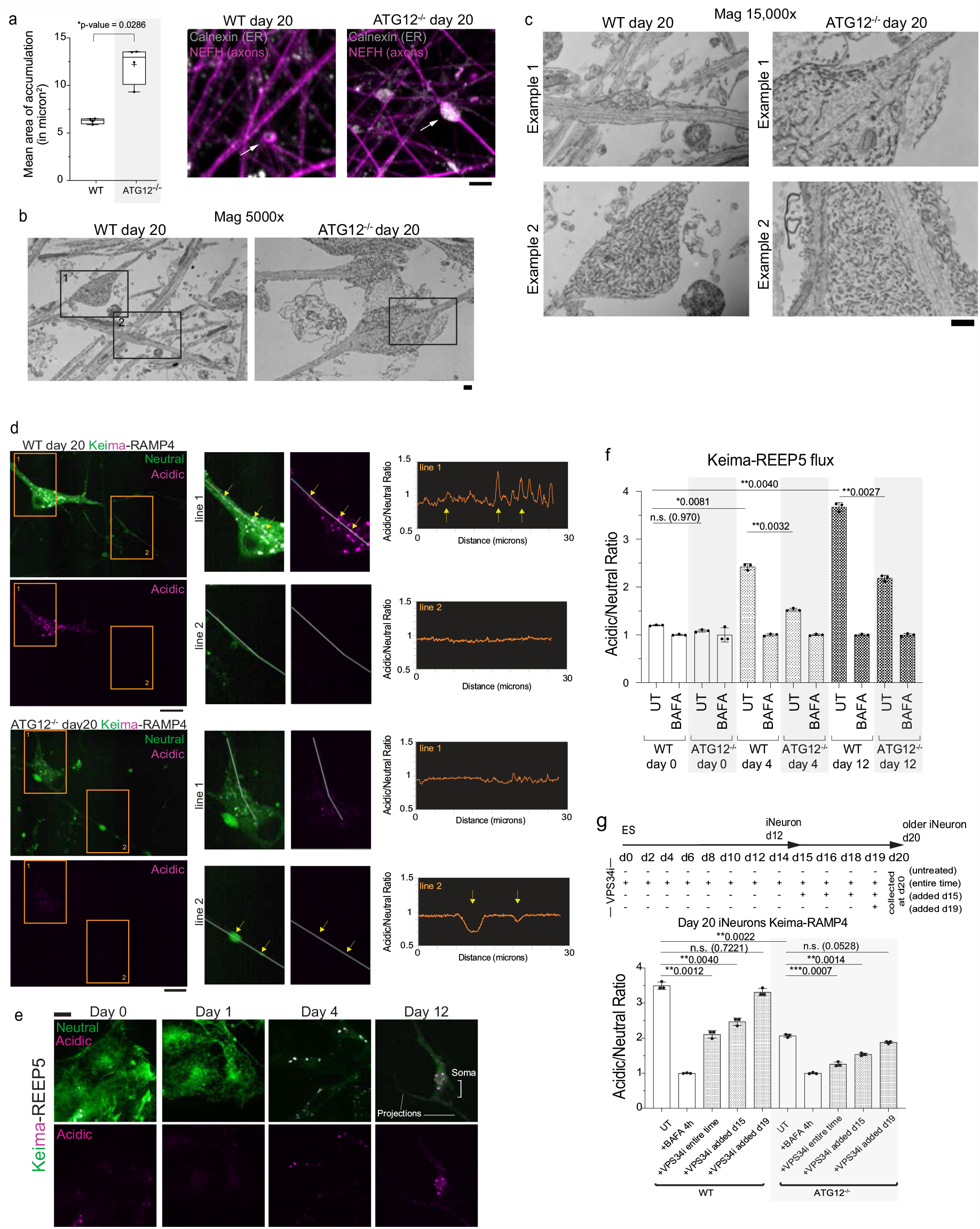
Autophagy-dependent clearance of ER in axons during iNeuron differentiation via ER-phagy receptors. **a**, As also represented in **Fig. 1b**, ATG12^-/-^ day 20 iNeurons were immunostained with α-NEFH and α-Calnexin to identify aberrant ER structures; here we compare these to WT day 20 iNeurons with the same immunostaining. Scale bar, 5 microns. Right panel, min-to-max box-and-whiskers plot representing median areas of ER accumulations. The line is at the median. + labels the mean. Four points for each condition represent the resulting median area from four independent differentiations. *, p<0.05; Mann-Whitney test. **b**,**c**, Scanning transmission EM of thin sections from WT and ATG12^-/-^ iNeuron cultures (day 20). Panel **b** shows low magnification image through multiple axons. Panel **c** displays high magnification images of example 1 and example 2 outlined in panel **b**. Scale bar, 500 nm. **d**, Live cells expressing Keima-RAMP4 in WT and ATG12^-/-^ day 20 iNeurons were imaged. Scale bar, 10 microns. Insets show the results of acidic/neutral ratiometric line scan analysis for somata or axons of WT (top panels) or ATG12^-/-^ iNeurons. **e**, Induction of ER-phagic flux during iNeuron differentiation. hESCs expressing Keima-REEP5 were differentiated to iNeurons and Keima signal imaged at day 0, 1, 4 and 12. Scale bar, 10 microns. **f**, Ratiometric analysis of WT or ATG12^-/-^ Keima-REEP5 flux was measured by flow cytometry at day 0, 4 and 12 of differentiation. The ratio of acidic to neutral Keima fluorescence was normalized to samples treated with BAFA (100 nM) for 4 h prior to analysis. Each measurement reflects biological triplicate measurements. **g**, Ongoing ER-phagic flux in day 15 iNeurons. WT or ATG12^-/-^ hESCs were differentiated in the presence or absence of VPS34i as indicated in the scheme. In some cases, VPS34i was added at day 19 or day 15 and Keima flux analyzed by flow cytometry, as in panel **f**. Each measurement reflects biological triplicate measurements. For **f**, and **g**, *, p<0.05; **, p<0.01; ***, p<0.001; n.s., not significant; Brown-Forsythe and Welch One-way ANOVA and Dunnett’s T3 multiple comparisons test.

**Extended Data Fig. 4.**
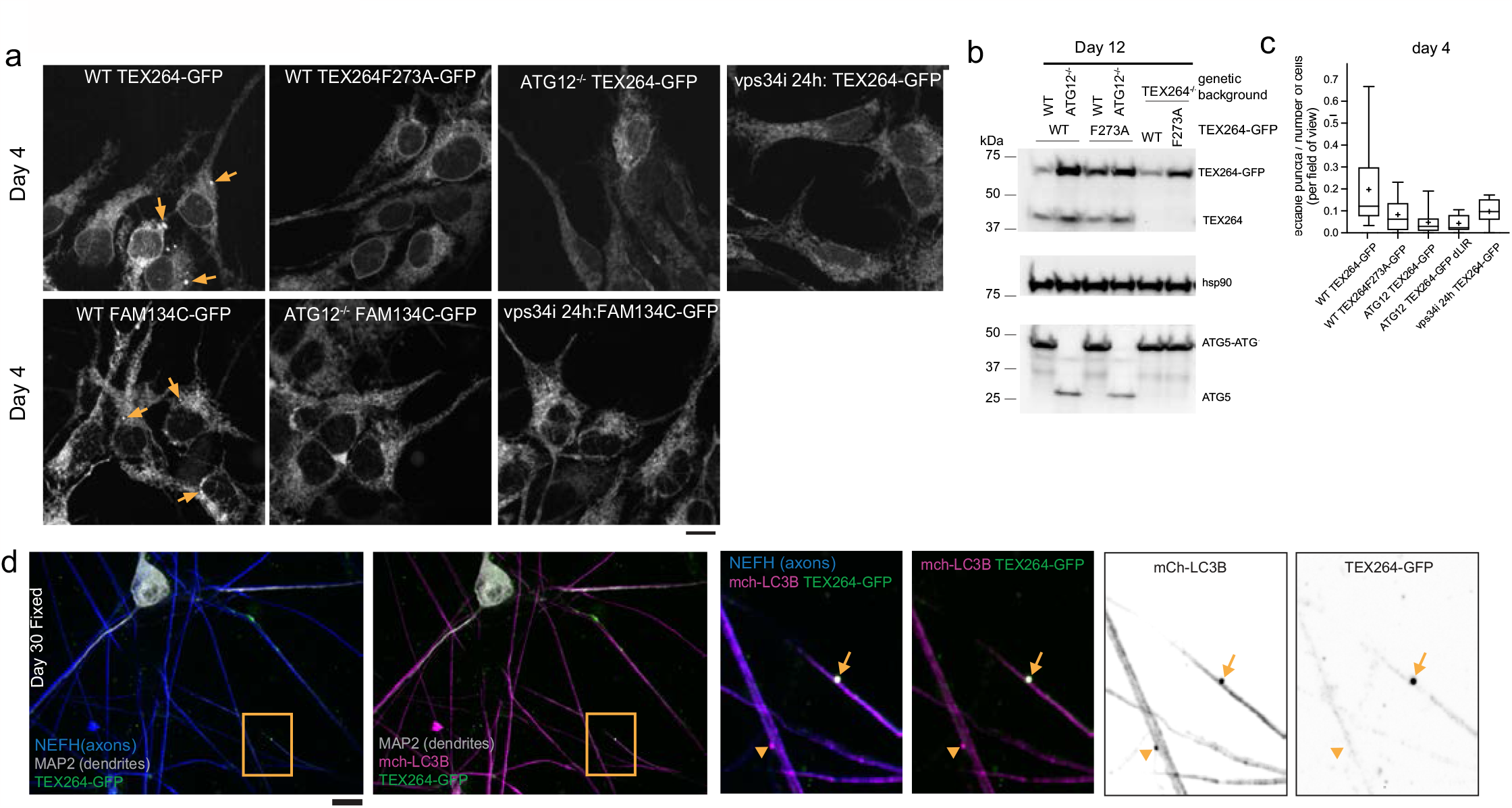
TEX264 and FAM134C puncta detection and tracking during iNeuron differentiation. **a-c**, TEX264-GFP, TEX264^F273A^-GFP, or FAM134C-GFP were expressed in WT or ATG12^-/-^ hESCs and cells imaged at day 4 of differentiation to iNeurons (panel **a**). In some experiments, VPS34i was added to WT cells for 24h prior to imaging. Arrows mark examples of ER-phagy receptor puncta. In panel **b**, expression of TEX264-GFP was verified by immunoblotting of iNeuron extracts using α-HSP90 as a loading control. In panel **c**, the number of TEX264-GFP puncta was quantified in day 4 iNeurons. **d** Day 30 iNeurons expressing TEX264-GFP and mCh-LC3B were immunostained with α-MAP2 to detect dendrites (white) and α-NEFH to mark axons (blue) and projections imaged by confocal microscopy. Insets show TEX264-GFP/mCh-LC3B-positive puncta (arrows) or mCh-LC3B-positive but TEX264-GFP-negative puncta (arrowheads) in axons. All scale bars, 10 microns.

**Extended Data Fig. 5.**
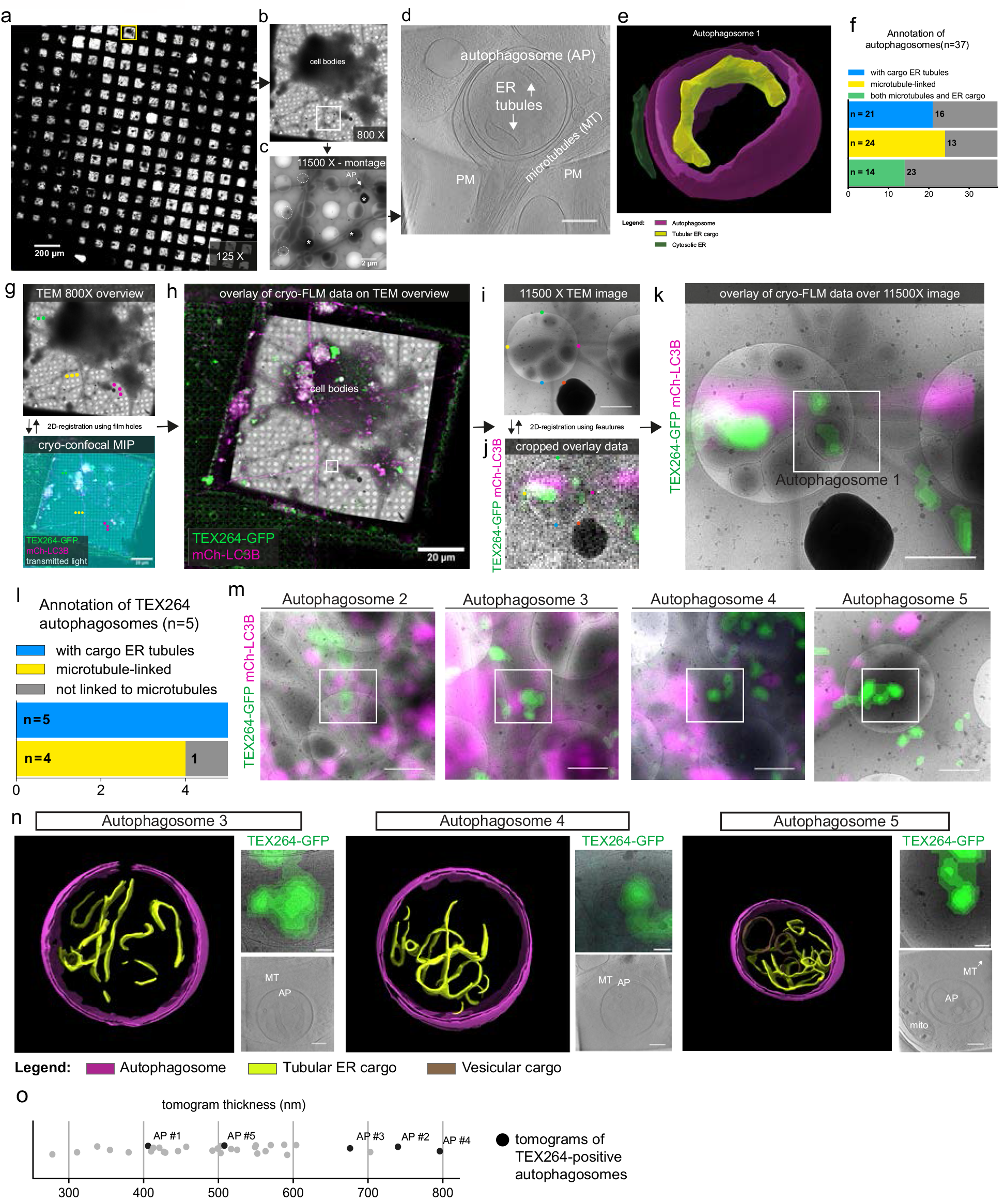
Analysis of autophagosomes in iNeurons by correlative cryo-ET. **a**,125X TEM image of the grid used in this study. The yellow square indicates the region shown in b. **b**, 800X TEM overview of a grid square. The darker area corresponds to the thicker iNeuron cell bodies; around it, many thinner projections corresponding to axon and dendrites can be observed. The white square indicates a particular area rich in neuronal projections, shown in c. **c**, 11500X TEM montage acquired to map a potentially interesting area. Extracellular vesicles are also present (white dashed circles). Pieces of ice are sitting on top of the frozen sample (asterisks). Autophagosome 1 (AP) was found in this area. **d**, Slice of the tomogram of autophagosome 1, denoised with Cryo-CARE. Scale bar, 200 nm. **e**, 3D rendering of autophagosome 1 with manually refined segmentations of a cytosolic ER tubule (dark green) and a portion of the tubular ER cargo (yellow). **f**, Cumulative barplot showing number of autophagosomes containing tubular ER cargo (n=21), linked to microtubules (n=24) or both (n=14). **g**, Example of the 2D-correlation workflow for autophagosome 1. Correlation of the 800X TEM overview with cryo-confocal data, represented by a single in-focus slice of the transmitted light stack (cyan) and maximum intensity projection (MIP) of fluorescence channels. The colored dots indicate the position of the 2 μm holes that were selected in both images for 2D-registration. After 2D registration, the TEM overview is transformed to match the fluorescence data. **h**, Overlay of the fluorescence data with the transformed TEM overview. The white square indicates the cropped area used for the second step of the correlation procedure. **i**, 11500X TEM image of the area around autophagosome 1. The colored dots indicate features of the image that were used for finer correlation with the cropped fluorescence data. **j**, Cropped fluorescence data corresponding to the square in g. The colored dots indicate the same correlation points shown in h. The fluorescence data is subsequently transformed to match the 11500X TEM image. **k**, Final overlay of the fluorescence data over the 11500X TEM image. The white square represents the tomogram position. **l**, Cumulative barplot showing number of TEX264-GFP positive autophagosomes that contain ER tubular cargo (n=5) and are linked to microtubules (n=4). **m**, 11500X TEM and fluorescence overlays for TEX264-GFP-positive autophagosomes 2, 3, 4 and 5. **n**, 3D segmentations, zoomed-in 11500X TEM images overlayed with GFP cryo-fluorescence signal and cryo-CARE denoised slices for autophagosomes 3, 4, and 5. All images are rotated right by 90 degrees compared to their respective full 11500X overlays in **m**. Autophagosome 3 and 4 are microtubule-linked, while autophagosome 3 is distant from the microtubules (MT). **o**, Plot showing thickness of all tomograms (n=32) analyzed in this work. Each dot represents a tomogram. Black dots indicate the tomograms corresponding to the 5 TEX264-GFP-positive autophagosomes (AP) shown in this study.

**Extended Data Fig. 6.**
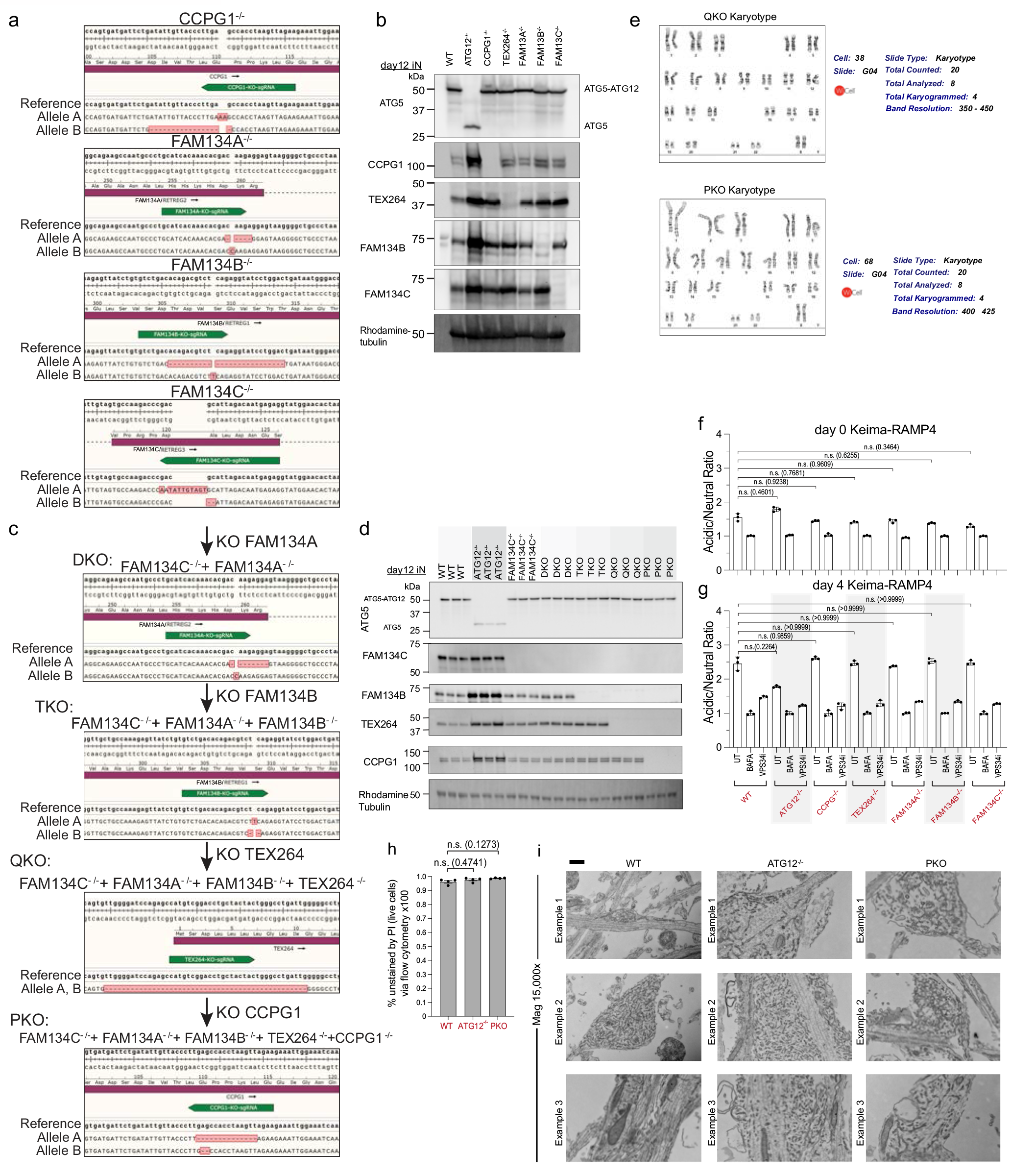
Generation of a genetic tool kit for functional analysis of ER-phagy receptors in iNeurons. **a**, MiSeq analysis of single ER-phagy receptor mutants in hESCs. The green highlights the target of the CRISPR gRNA. The sequence of the major MiSeq output is indicated for each allele. **b**, Immunoblot validation of targets knock-out clones at day 12 of differentiation. Cell extracts were subjected to immunoblotting with the indicated antibodies, employing a Rhodamine-labeled α-tubulin as loading controls. **c**, MiSeq analysis of combinatorial ER-phagy receptor mutants in hESCs, as performed for the single knockouts in a. **d**, Immunoblot validation of targets in combinatorial knock-out clones at day 12. Cell extracts were subjected to immunoblotting as in **b. e**, Karyotype analysis of QKO and PKO hESCs revealed no detectable alterations in chromosome number. **f**, Ratiometric flow cytometry analysis of Keima-RAMP4 flux was measured in WT, ATG12^-/-^, or the indicated ER-phagy receptor knock-out ES cells (day 0 of differentiation). The ratio of acidic to neutral Keima fluorescence was normalized to samples treated with BAFA (100 nM) for 4 h prior to analysis, and where indicated, cells were cultured with VPS34i prior to analysis. Each measurement (represented by a point) reflects a biological triplicate sample. n.s. not significant. Error bars represent SD. **g**, As in panel **f**, but at day 4 of differentiation to iNeurons. For **f**, and **g**, n.s., not significant; Brown-Forsythe and Welch One-way ANOVA and Dunnett’s T3 multiple comparisons test. **h**, Day 12 iNeurons treated Propidium iodine staining and were analyzed via flow cytometry. The same gating strategy for live cells was applied to all genotypes, as was done in all Keima flux experiments. Percent of live cells (not stained with PI) is displayed. N=3 biological replicates; n.s. not significant; ; Mann-Whitney test. **i**, Examples of enlarged axonal structures from WT, ATG12^-/-^ and PKO day 30 iNeurons containing dense tubular ER, as visualized by TEM. Some of these examples were also shown in **Extended Data Fig. 1c** to compare only WT and ATG12^-/-^. Three independent examples are shown. Scale bar, 500nm.

**Extended Data Fig. 7.**
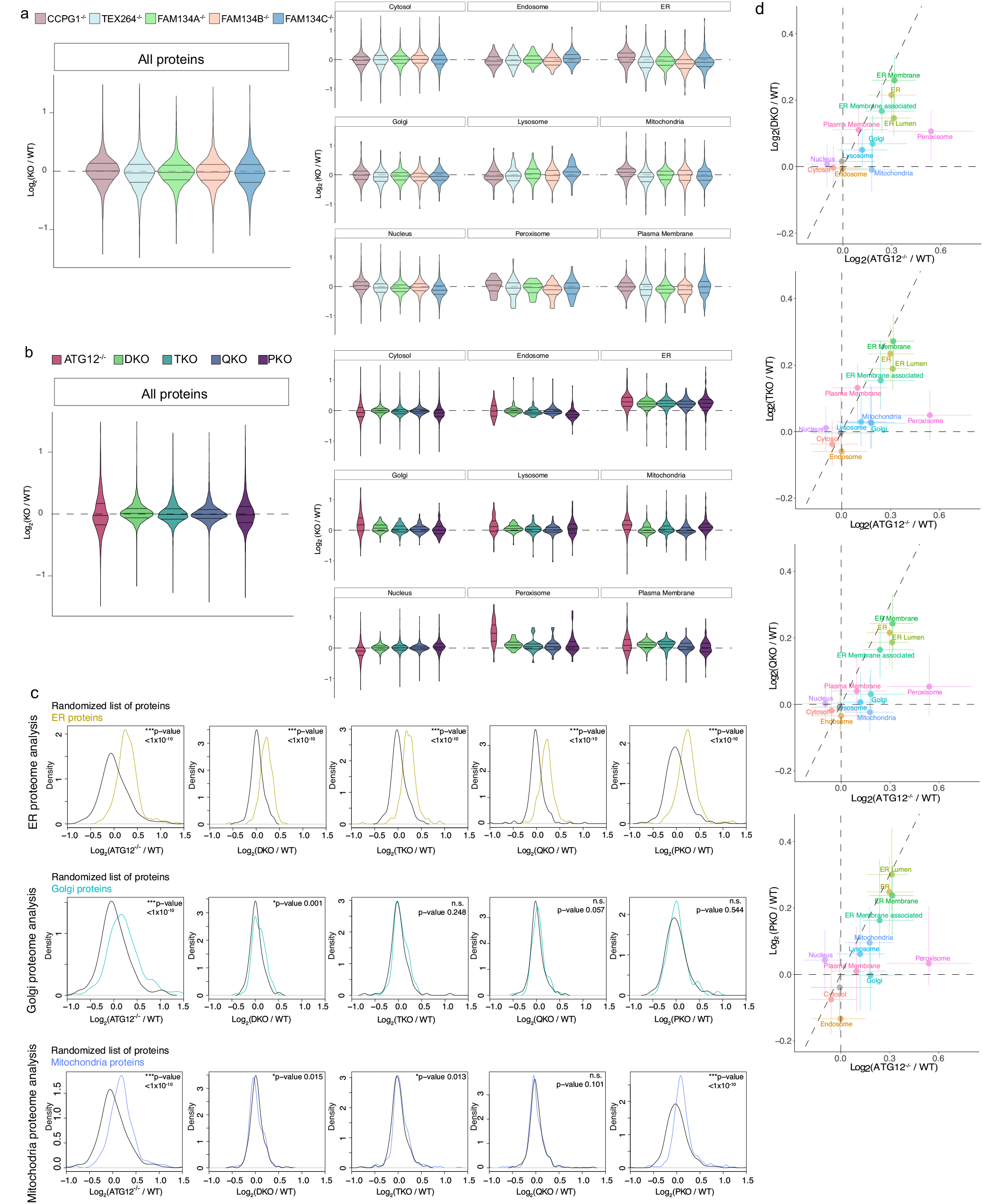
Combinatorial analysis of ER remodeling via ER-phagy receptors during neurogenesis in vitro. **a**, Violin plots for changes in individual organelle abundance in the indicated single ER-phagy knockout iNeurons (day 12). **b**, Violin plots for changes in individual organelle abundance in the indicated combinatorial ER-phagy knock-out iNeurons (day 12). **c**, Log_2_FC (mutant/WT) distributions of ER proteins (top), Golgi proteins (middle), or mitochondria proteins (bottom) compared to randomized selections of the same number of proteins (100 iterations). p-values for each comparison are calculated with a Kolmogorov-Smirnov Test (two-sided). The ER protein section of the figure (top panel) is reproduced from **Fig. 6c. d**, Correlation plots for changes in organelle abundance (Log_2_FC) comparing DKO, TKO, QKO and PKO log_2_FCs from WT individually with ATG12^-/-^ log_2_FCs from WT.

**Extended Data Fig. 8.**
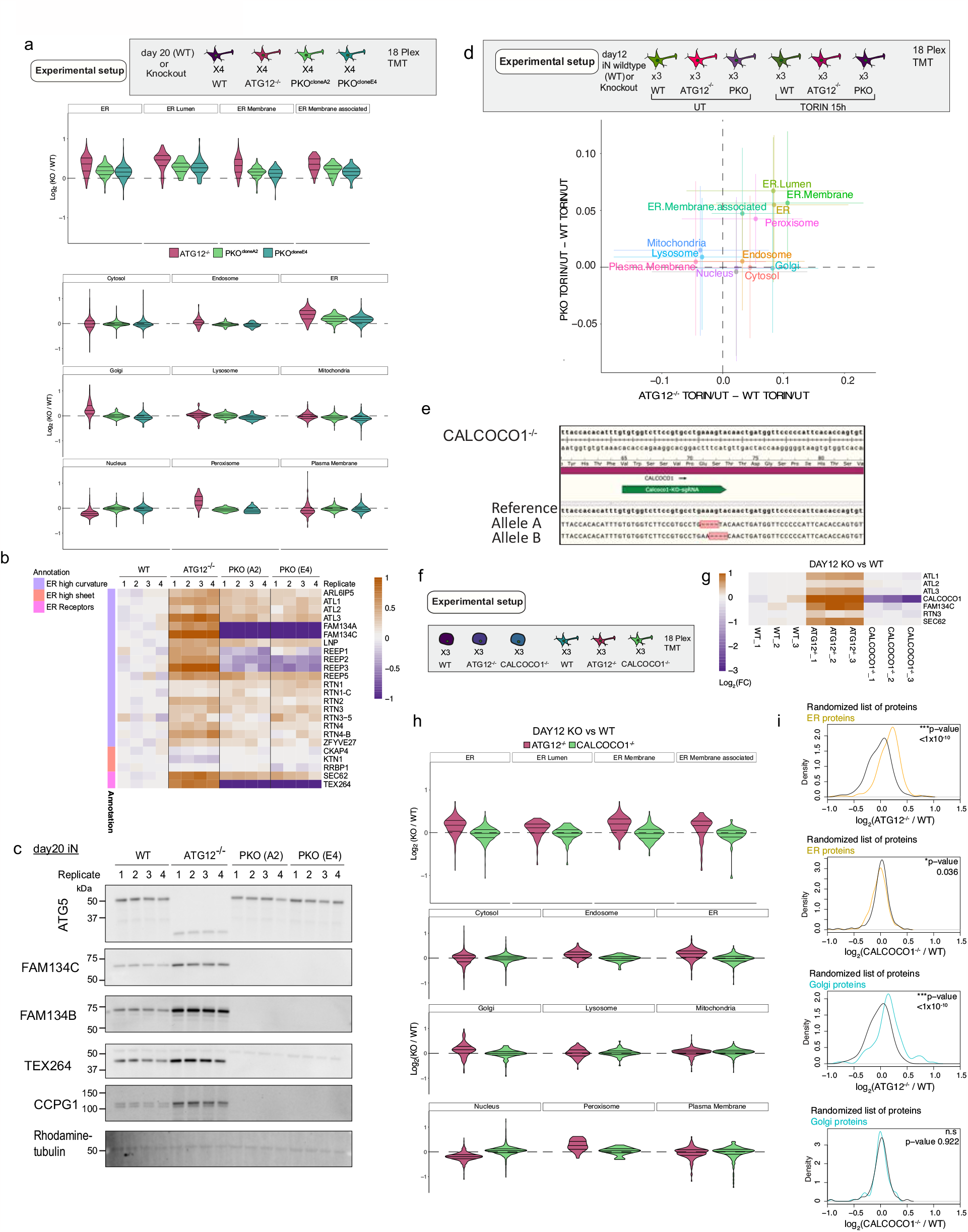
Analysis of ER-proteome remodeling in PKO and CALCOCO1^*-/-*^ iNeurons. **a**, Proteomic analysis of two PKO clones (A2 and E4) in parallel with ATG12^-/-^ and WT iNeurons (day 20). The upper panel provides a schematic of the TMT multiplex approach employed. n=4 biological replicates. Middle panel displays Log_2_FC for ER protein and selected ER protein categories. Lower panel displays Log_2_FC for the indicated organelles. **b**, Heat map for Log2FC values for the indicated proteins from the experiment in panel **a. c**, Immunoblot validating deletion of FAM134B, FAM134C, TEXT264, and CCPG1 in both clone A2 and E4 for the PKO mutant. **d**, Modulation of iNeuron proteome in response to inhibition of MTOR with Torin1 (100 nM,15 h). Upper panel shows a schematic of the experimental set-up employing TMT based proteomics to quantity alterations in the proteome if WT or ATG12^-/-^, PKO iNeurons. Lower panel: Correlation plots comparing the effect of Torin1 on organelles of ATG12^-/-^ cells relative to WT cells and PKO cells relative to WT cells. **e**, MiSeq analysis of a CALCOCO1-/-H9 hESC clone, showing the position of the gRNA used for CRISPR-Cas9 deletion, and the position of out of frame deletions in the two alleles of CALCOCO1. **f**, Schematic showing the TMT total proteome strategy for analysis of the effect of CALCOCO1 deletion on organelle abundance in ES cells and day 12 iNeurons (n=3 biological replicates). ATG12^-/-^ cells were included as a positive control. **g**, Heat map of Log_2_FC values for selected proteins from the TMT experiment outlined in panel **f**, which also demonstrates loss of the CALCOCO1 protein in the CALCOCO1^-/-^ iNeurons. **h**, Violin plots (Log_2_FC) for the indicated organelles (top panel) and selected classes of ER proteins (lower panel) for ATG12^-/-^ and CALCOCO1^-/-^ iNeurons (day 12). Loss of CALCOCO1 does not affect the abundance of any of the organelles tested. **i**, Comparison of Log_2_FC (mutant/WT) distributions of ER proteins (top) or Golgi proteins (bottom) to distributions of randomized selections of the same number of proteins (100 iterations). p-values for comparisons were calculated with a Kolmogorov-Smirnov Test (two-sided).

**Extended Data Fig. 9.**
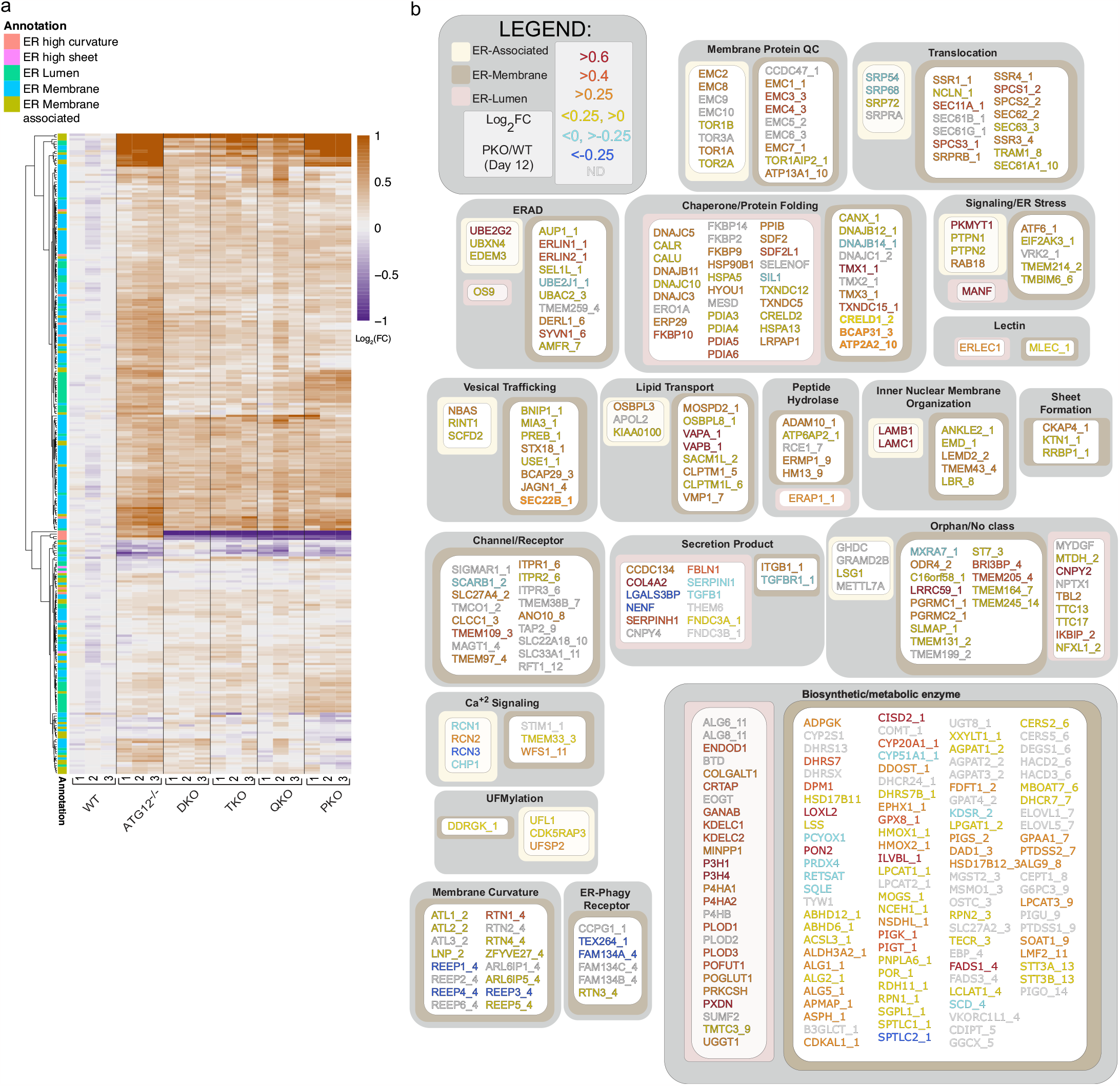
Overview of ER proteome remodeling via ER-phagy receptors during neurogenesis in vitro. **a**, Changes in the abundance (Log_2_FC) of the ER proteome (267 detected proteins) during conversion in ATG12^-/-^ or combinatorial ER-phagy receptor knockout iNeurons (day 12) are shown in as heatmaps. Annotations of the type of ER protein are indicated by the relevant colors. **b**, Landscape of the ER proteome and the effect of deletion of five ER-phagy receptors (FAM134A/B/C, TEX263 and CCPG1) on accumulation of individual proteins. The ER proteome (359 proteins, **Supplementary Data Table 1**) is organized into functional modules and protein attributes (involved in ER membrane curvature, ER-associated, ER-membrane, ER-Lumen or ER-phagy receptor) are indicated by the respective outline box color (see inset legend). For proteins with transmembrane segments, the number of segments is indicated after the protein name (_1, _2, etc) based on data in Uniprot. The text of each protein name is colored based on day12 PKO vs WT Log_2_FC (see inset legend). (**Supplementary Data Table 3**).

**Extended Data Fig. 10.**
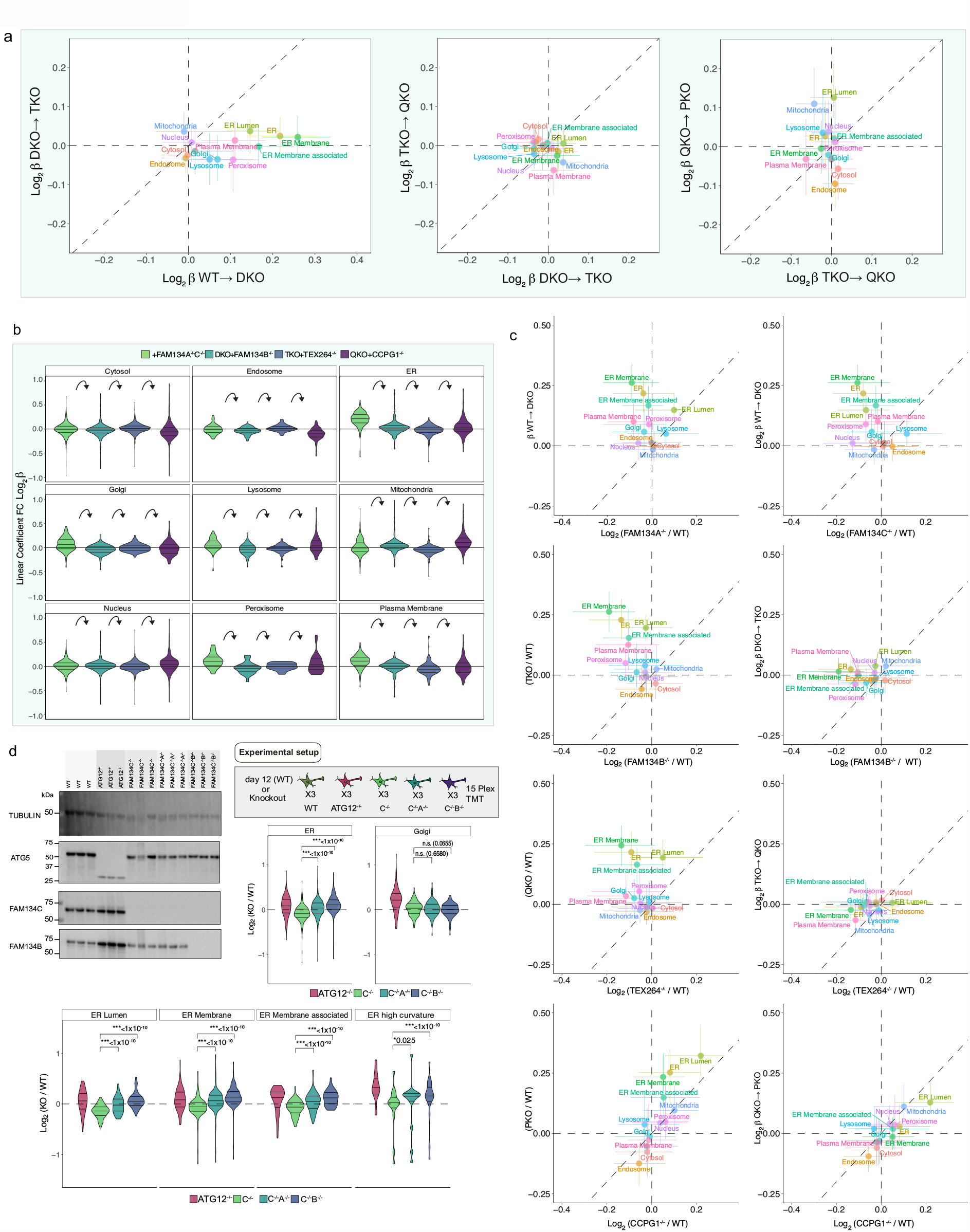
Application of a linear model for alterations in ER proteome abundance in sequential ER-phagy receptor knockout cells during iNeuron differentiation. **a**, Effect of sequential ER-phagy receptor deletion on the β coefficient values for individual organelles measured by quantitative proteomics in day 12 iNeurons. **b**, Violin plots reflecting changes in β coefficient values for individual organelles measured by quantitative proteomics in day 12 iNeurons. Curved arrows reflect sequential removal of the indicated ER-phagy receptor. **c**, Correlation plots for the indicated β coefficient or Log_2_FC plots comparing organelle abundance for combinatorial or single ER-phagy deletion iNeurons. **d**, Experiment characterizing FAM134C^-/-^, FAM134A/C^-/-^, and FAM134B/C^-/-^ day 12 iNeurons with ATG12^-/-^ iNeurons included as a control. Top left, Immunoblot of extracts from iNeurons of the indicated genotypes (n = 3) were probed with α-ATG5, α-FAM134B, α-FAM134C, or α-tubulin. ATG12^-/-^ cells display loss of the ATG12-ATG5 conjugate as observed with α-ATG. Top right, experimental scheme for multiplexed total proteome analysis of FAM134C^-/-^, FAM134A/C^-/-^, and FAM134B/C^-/-^ iNeurons and ER proteome specific violin plots derived from this analysis. Bottom, organelle-specific violin plots from the experiment. p-values for comparisons between violins were calculated using paired Wilcoxon tests.

**Extended Data Fig. 11.**
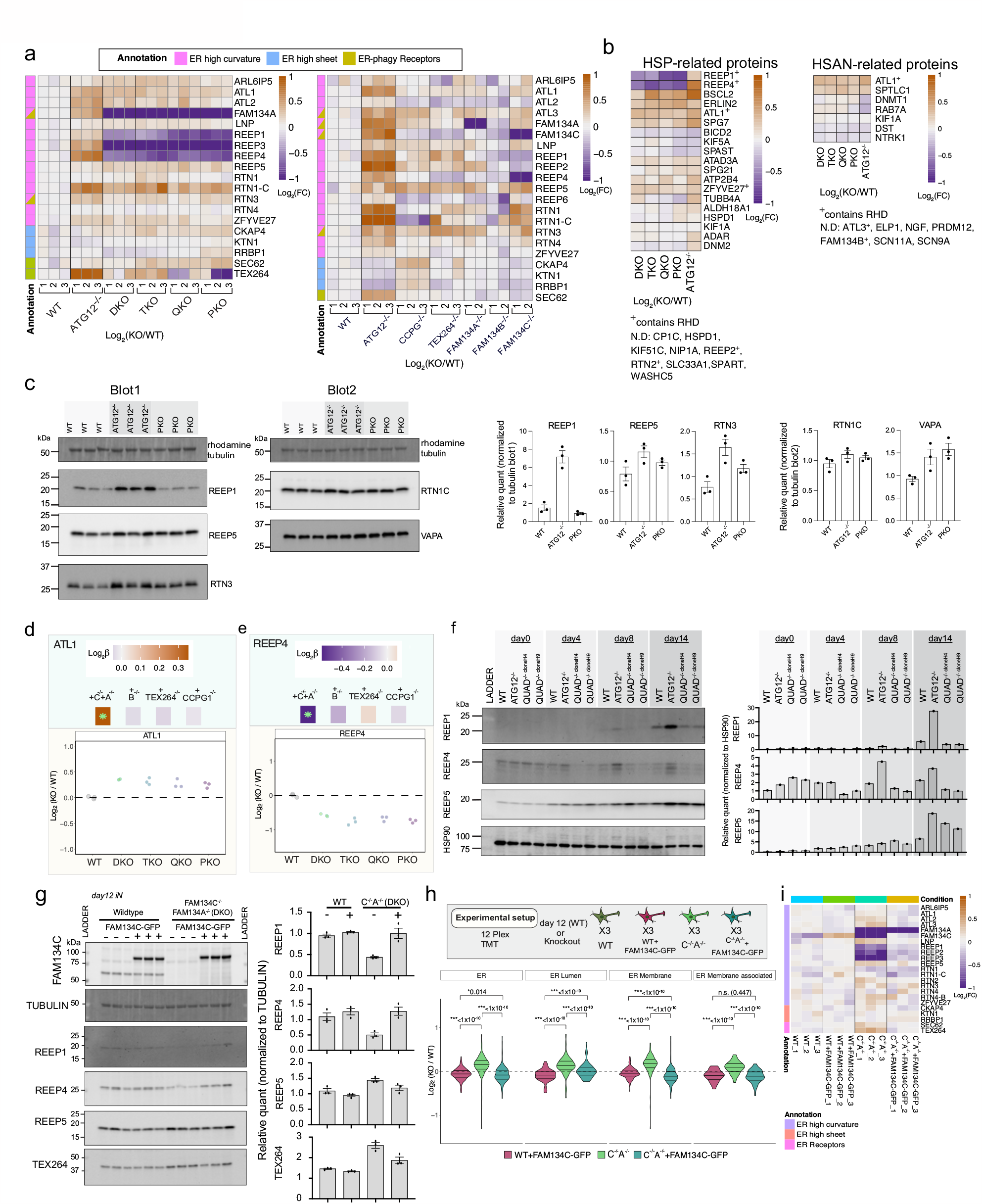
Differential regulation of ER membrane shaping and disease-linked proteins upon loss of ER-phagy receptors. **a**, Heat maps displaying Log_2_ FC values for selected ER-shaping proteins for the indicated iNeuron genotypes. **b**, Heat maps displaying Log_2_ FC values for Hereditary Spastic Parapelgia (HSP)-linked proteins or Hereditary Sensory and Autonomic Neuropathy (HSAN)-linked proteins for the indicated iNeuron genotypes. **c**, Immunoblot of extracts from WT, ATG12^-/-^, or PKO day 12 iNeurons were probed with α-REEP1, α-REEP5, α-RTN3 or α-tubulin as a loading control (blot1) or were probed with α-RTN1C, α-VAPA or α-tubulin as a loading control (blot2). The relative levels of proteins were quantified (right). **d-e**, Further examples ER shaping proteins that accumulate (**d**) or decrease (**e**) with additional ER-phagy receptor knockout. **f**, Immunoblot of cell extracts isolated from the indicated time point during differentiation, using α-HSP90 as a loading control. The relative levels of proteins were quantified (right). **g**, Immunoblot of extracts from WT or FAM134A/C^-/-^ (DKO) iNeurons with or without expression of FAM134C-GFP using a PiggyBac vector (n = 3). Blots were probed with α-TEX264, α REEP1, α-REEP4 or α-tubulin as a loading control. The relative levels of proteins were quantified in the lower panel. **h**, Experimental scheme for multiplexed total proteome analysis of WT or FAM134A/C^-/-^ with or without expression of FAM134C-GFP iNeurons (n = 3) and ER protein-specific violin plots derived from this analysis. p-values for comparisons between violins were calculated using paired Wilcoxon tests. **i**, Heat map displaying Log_2_ FC values for selected ER-shaping proteins for the indicated iNeuron genotypes.

**Extended Data Fig. 12.**
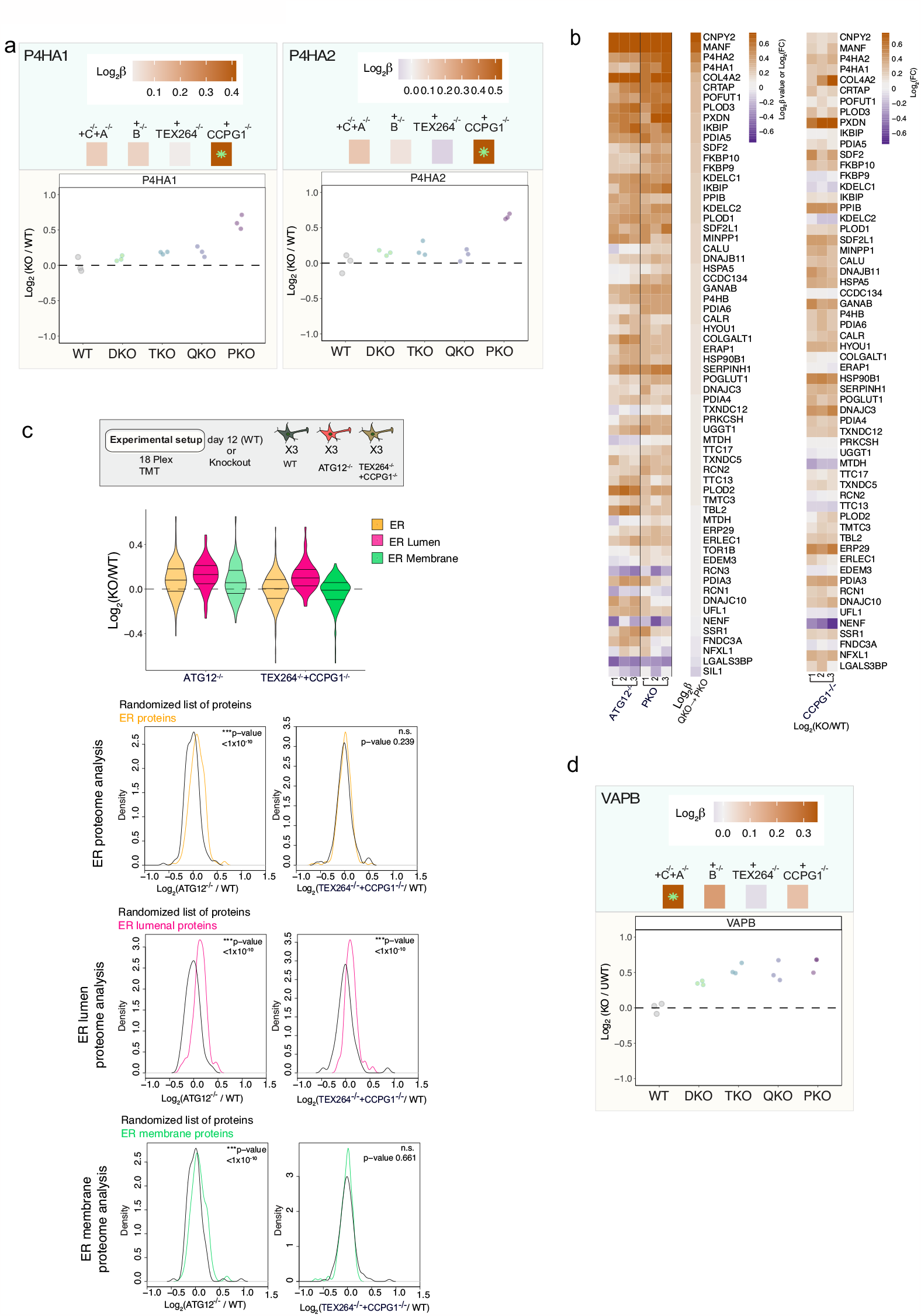
Differential regulation of ER membrane shaping proteins upon loss of ER-phagy receptors. **a**, ER lumenal proteins that accumulate with additional ER-phagy receptor knockout. Top panels are β coefficient values and lower panels are Log_2_FC; green asterisks in β coefficient for single protein heat maps indicate significant change (adjusted p-value < 0.05) in β coefficient. **b**, ER lumenal protein heatmaps reflecting the change in abundance (Log_2_FC) for deletion of ATG12 or PKO, reflecting β coefficient values for QKO to PKO, (left panel) or reflecting the change in abundance (Log_2_FC) for single deletion of CCPG1 (right panel). **g**, Experimental scheme for multiplexed total proteome analysis of WT, ATG12^-/-^ or CCPG1^-/-^/TEX264^-/-^ iNeurons (n = 3) and violin plots for total ER, ER-lumen, and ER-membrane proteins derived from this analysis. **c**, Violin plots from a TMT 18plex experiment comparing WT, ATG12^-/-^, and another ER-phagy receptor allelic combination (TEX264^-/-^ +CCPG1^-/-^) with complementary comparisons of Log_2_FC (mutant/WT) distributions of ER proteins (top), ER lumenal proteins (middle), or ER membrane proteins (bottom) to distributions of randomized selections of the same number of proteins (100 iterations). p-values for each comparison are calculated with a Kolmogorov-Smirnov Test (two-sided) test. Together these reflect accumulation of the ER, ER lumen and ER membrane for Log2FC (ATG12^-/-^/WT) but only the ER lumenal proteome accumulates for Log_2_FC (TEX264^-/-^ +CCPG1^-/-^ /WT). **d**, ER contact site protein VAPB accumulates with additional ER-phagy receptor knockout. Plot is annotated as in panel **a**.

## Supplemental Tables

**Supplementary Data Table 1**. ER protein designation table. The ER proteome (359 proteins) is organized into functional modules and protein attributes (involved in ER membrane curvature, ER-associated, ER-membrane, ER-Lumen or ER-phagy receptor). For proteins with transmembrane segments, the number of segments is indicated after the protein name (_1, _2, etc) based on data in Uniprot. Relevant to assigning ER designations in all other Supplementary Data tables. In .xlsx format

**Supplementary Data Table 2**. Total proteome analysis for ESC conversion to iNeurons, WT day 0 to day 12. Relevant to Fig. 1a,b and Extended Data Fig. 1b. In .xlsx format. Details of the experiment underlying this data are provided in a tab in the table.

**Supplementary Data Table 3**. Total proteome analysis for WT, ATG12^-/-^, DKO, TKO, QKO, PKO day12 iNeurons. ATG12^-/-^ vs. WT relevant to Fig. 1c,d,e and Extended Data Fig. 1a. ATG12^-/-^, DKO, TKO, QKO, PKO versus WT relevant to Fig. 6b-g, 7a-c,e,f; Extended Data Fig. 7b-d, 9, 10a-c, 11a,b,d, 12a,b,d. In .xlsx format. Details of the experiment underlying this data are provided in a tab in the table.

**Supplementary Data Table 4**. Total proteome analysis for WT, ATG12^-/-^, PKO day 0 ESC and day12 iNeurons. Relevant to Extended Data Fig. 2a. In .xlsx format. Details of the experiment underlying this data are provided in a tab in the table.

**Supplementary Data Table 5**. Total proteome analysis for WT, ATG12^-/-^, CCPG1^-/-^, TEX264^-/-^, FAM134A^-/-^, FAM134B^-/-^, and FAM134C^-/-^ day12 iNeurons. Relevant to Fig 6a and Extended Data Fig. 7a, 10c, 11a, 12b. In .xlsx format. Details of the experiment underlying this data are provided in a tab in the table.

**Supplementary Data Table 6**. Total proteome analysis for WT, ATG12^-/-^, PKO^cloneA2^, PKO^cloneE4^ day20 iNeurons. Relevant to Extended Data Fig. 8a-c. In .xlsx format. Details of the experiment underlying this data are provided in a tab in the table.

**Supplementary Data Table 7**. Total proteome analysis for WT, ATG12^-/-^, PKO day12 iNeurons with and without Torin1 inhibitor. Relevant to Extended Data Fig. 8d. In .xlsx format. Details of the experiment underlying this data are provided in a tab in the table.

**Supplementary Data Table 8**. Total proteome analysis for WT, ATG12^-/-^, CALCOCO1^-/-^ day0 and day12 iNeurons. Relevant to Extended Data Fig. 8e-i. In .xlsx format. Details of the experiment underlying this data are provided in a tab in the table.

**Supplementary Data Table 9**. Total proteome analysis for WT, ATG12^-/-^, FAM134C^-/-^, FAM134CA^-/-^(DKO), and FAM134CB^-/-^ day12 iNeurons. Relevant to Extended Data Fig. 10d. In .xlsx format. Details of the experiment underlying this data are provided in a tab in the table.

**Supplementary Data Table 10**. Total proteome analysis for WT, WT+FAM134C-GFP, FAM134CA^-/-^(DKO), and FAM134CA^-/-^(DKO)+FAM134C-GFP day12 iNeurons. Relevant to Extended Data Fig. 10g-i. In .xlsx format. Details of the experiment underlying this data are provided in a tab in the table.

**Supplementary Data Table 11**. Total proteome analysis for WT, ATG12^-/-^, TEX264^-/-^+CCPG1^-/-^ day12 iNeurons. Relevant to Extended Data Fig. 11c. In .xlsx format. Details of the experiment underlying this data are provided in a tab in the table.

## Supplemental Movies

**Supplemental Movie 1, related to Fig. 3**. An additional example of day 30 WT iNeurons expressing TEX264-GFP (green) and mCh-LC3B (red) imaged every 30s. Image sequence played at 2 frames per second. Circles indicate co-trafficking of TEX264-GFP/mCh-LC3B positive puncta. Scale bar, 10 microns.

**Supplemental Movie 2, related to Fig. 3**. Day 30 WT iNeurons expressing FAM134C-GFP (green) and mCh-LC3B (red) imaged every 30s. This example movie includes the cropped region represented in Fig. 2j. Image sequence played at 2 frames per second. Circles indicate co-trafficking FAM134C-GFP/mCh-LC3B positive puncta. Scale bar, 10 microns.

**Supplemental Movie 3, related to Fig. 3**. A region of a day 30 WT iNeuron expressing TEX264-GFP (green) and mCh-LC3B (red) imaged every 20s. This example movie is the region represented in Fig. 2k where a TEX264-GFP/mCh-LC3B positive puncta leaves the dilated axonal region. Image sequence played at 2 frames per second. Scale bar, 5 microns.

**Supplemental Movie 4, related to Fig. 4**. The tomogram series for autophagosome 1 captured *in situ* by cryo-ET.

## METHODS

Protocols associated with this work can be found on protocols.io with the following DOI: dx.doi.org/10.17504/protocols.io.81wgbx13nlpk/v2

### Reagents

The following chemicals, peptides, and recombinant proteins were used: DAPI Thermo Fisher Scientific (D1306); TMTpro™ 16plex Label Reagent Set Thermo Scientific (A44520); Q5 Hot Start High-Fidelity DNA Polymerase New England BioLabs (M0493); Gateway LR Clonase II Enzyme Mix Thermo (11791020); NEBuilder HiFi DNA Assembly Master Mix (E2621s); MiSeq Reagent Nano Kit v2 (300 cycles) Illumina (MS-103-1001); Bafilomycin A1 Cayman Chemical (88899-55-2); Sar405 Selective ATP-competitive inhibitor of Vps34 Apexbio (A8883); DAPI (4’,6-Diamidino-2-Phenylindole, Dihydrochloride) Thermo Fisher Scientific (D1306); 16% Paraformaldehyde, Electron-Microscopy Grade Electron Microscopy Science (15710), PhosSTOP Sigma-Aldrich (T10282); Protease inhibitor cocktail Roche (4906845001); TCEP Gold Biotechnology, Formic Acid Sigma-Aldrich (94318); Trypsin Promega (V511C); Lys-C Wako Chemicals (129-02541); Urea Sigma (U5378); EPPS Sigma-Aldrich (E9502); 2-Chloroacetamide Sigma-Aldrich C0267; Trypan Blue Stain Thermo Fisher Scientific Wako Chemicals (129-02541w); Urea Sigma (U5378); EPPS Sigma-Aldrich (E9502); 2-Chloroacetamide Sigma-Aldrich (C0267); Empore SPE Disks C18 3M Sigma-Aldrich (66883-U); GeneArt Precision gRNA Synthesis Kit Thermo Fisher Scientific (A29377); 12 Well glass bottom plate with high performance #1.5 cover glass Cellvis (P12-1.5H-N); Nunc Cell-Culture Nunclon Delta Treated 6-well Thermo Fisher Scientific (140685); Nunc Cell-Culture Nunclon Delta Treated 12-well Thermo Fisher Scientific (150628); 100×21mm Dish, Nunclon Delta Thermo Fisher Scientific (172931); Corning Matrigel Matrix, Growth Factor Reduced Corning (354230); DMEM/F12 Thermo Fisher Scientific (11330057); Neurobasal Thermo Fisher Scientific (21103049); Non Essential Amino Acids Life Technologies (11140050); GlutaMax Life Technologies (35050061); N-2 Supplement Thermo Fisher Scientific (17502048); Neurotrophin-3 (NT-3) Peprotech (450-03); Brain-derived neurotrophic factor (BDNF) Peprotech (450-02); B27 Thermo Fisher Scientific (17504001); Y-27632 Dihydrochloride (ROCK inhibitor) PeproTech (1293823); Cultrex 3D Culture Matrix Laminin I R&D Systems (3446-005-01); Accutase StemCell (7920); FGF3 In-house (N/A); Insulin Human Sigma-Aldrich (I9278-5ML); TGF-beta PeproTech (100-21C); holo-Transferrin human Sigma-Aldrich (T0665); Sodium Bicarbonate Sigma-Aldrich (S5761-500G); Sodium selenite Sigma-Aldrich (S5261-10G); Doxycycline Sigma-Aldrich (D9891); Recombinant SpCas9 ^51^; Hygromycin B Thermo Fisher Scientific (10687010); UltraPure 0.5M EDTA, pH 8.0 Thermo Fisher Scientific (15575020); GlutaMAX Thermo Fisher Scientific (35050061); Dulbecco’s MEM (DMEM), high glucose, pyruvate GIBCO / Invitrogen (11995); Lipofectamine 3000 Invitrogen (L3000008); Click-iT Plus TUNEL assay (Invitrogen C10617, with Alexa Fluor 488); Tunicamycin Cell Signaling (12819S); RNAeasy Qiagen kit (Qiagen 74104); Qiashredder columns (Qiagen 79654); DNAseI (Thermo EN0521); oligo dT_20_ primers (Invitrogen 79654); dNTPs (NEB N0447L)

### Plasmids

Plasmids constructed for and used in this manuscript will be available at Addgene upon final publication. These include pAC150-Keima-RAMP4 (this paper, Addgene 201929); pAC150-Keima-VAPA (this paper, Addgene in process); pAC150-Keima-REEP5 (this paper, Addgene 201928); pAC150-FAM134C-GFP (this paper, Addgene 201932); pAC150-TEX264-GFP (this paper, Addgene 201931); pAC150-TEX264(deltaLIR, F273A)-GFP (this paper, Addgene 201930), pHAGE-FAM134C-GFP (this paper, Addgene 201927); pHAGE-TEX264-GFP (Addgene 201925 ^10^); pHAGE-TEX264(deltaLIR,F273A)-GFP (Addgene 201926 ^10^); pHAGEmCherry-LC3B (Addgene 201924 ^10^).

### Cell Culture

Human embryonic stem cells (hESC, H9, WiCell Institute, WA9, RRID: CVCL_9773) were cultured in E8 medium on Matrigel coated plates as described^26^. Cells were split when they reached 80% confluency (every 2-4 days) using 0.5 mM EDTA in 1× DPBS (Thermo Fisher Scientific).

### Neural differentiation of AAVS1-TRE3G-NGN2 pluripotent stem cells

TRE3G-NGN2 was integrated into the AAVS site of the hESCs and iPSCs as previously described^52^. To start differentiation to induced neurons (iNeurons) from stem cells (day 0), cells were plated at 2×10^5^ cells/ml onto Matrigel coated plates into ND1 medium (DMEM/F12, 1X N2 (thermo), human Brain-derived neurotrophic factor BDNF (Brain derived neurotrophic factor) (10 ng/ml, PeproTech), human Neurotrophin-3 NT3 (10 ng/ml, PeproTech), 1X NEAA (Non-essential amino acids), Human Laminin(0.2 ug/ml) and Doxycycline (2 mg/ml)) also containing Y27632 (rock inhibitor-10 mM)). The media was replaced with ND1 without Y27632 the next day. The following day media was replaced with ND2 (Neurobasal medium, 1X B27, 1X Glutamax, BDNF (10 ng/ml), NT3 (10 ng/ml) and doxycycline at 2 mg/ml. On Days 4 and 6, 50% of the media was changed with fresh ND2. On any day in the Day 4-7 range, cells were replated at 4×10^5^ cells/well in ND2 medium with Y27632. The media was replaced the next day with fresh ND2 (without Y27632) Every other day 50% of the media was changed with ND2. At Day9 and onwards doxycycline was removed from the ND2 mixture. iNeurons were fed every other day with 50% media change until the experimental day (Day12 of differentiation unless otherwise noted).

### Molecular Cloning

Plasmids were made using either Gateway technology (Thermo) or via Gibson assembly (New England biolabs) in pHAGE backbone (for lentivirus transduction) or in the pAC150 piggy Bac backbone (for stable hESC generation). Entry clones from the human orfeome collection version 8 were obtained and cloned via LR cloning into various destination expression vectors.

### Lentivirus generation and Viral transduction of induced neurons

Lentiviral vectors were packaged in HEK293T (ATCC, CRL-1573, RRID: CVCL_0045) by cotransfection of pPAX2, pMD2 and the vector of interest in a 4:2:1 ratio using Lipofectamine 3000. One day after transfection media was changed to ND2 (no DOX) and then the following day virus containing supernatant was collected, filtered through a .22 micron syringe filter and frozen at -80°C. hESCs were differentiated to neurons as described above. At day 11 (two days after Dox removal) the iNeurons were transduced. iNeurons were imaged one day after transduction or at any following day (experimental day noted in each figure).

### Electroporation and selection of stable hESC populations with PiggyBac vectors

PiggyBac plasmids freshly maxi prepped at high concentrations were electroporated into hESCs using the 10μL Neon ThermoFisher kit and ThermoFisher Neon Electroporator. 1.5 μg of pAC150 piggy Bac vectors for ER proteins (Keima-RAMP4, TEX264-GFP, FAM134C-GFP and 1 μg of pCMV-hyPBase hyperactive piggyBac vector. 2×10^5^ cells 10μl buffer R were used for each electroporation. Program 13 was used from the optimization tab for electroporation parameter (Voltage: 1100. Pulse width: 20 Pulse number: 2). We plated the electroporated ESCs into Matrigel coated plates containing E8 with Y27632 (rock inhibitor-10 mM) and cells were placed in a low O_2_ incubator for two to four days. After four days with regular E8 media changes daily (or when cells reach 80 percent confluency) the cells were split into selection media (E8 with Y27632 and 50μg/mL hygromycin B). Cell were grown in selection medium for 7-10 days until there was no longer any cell death. Cells were further selected to obtain a fluorophore-positive population via flow cytometry with Sony Biotechnology (SH800S) Cell Sorter.

### Gene Editing

Gene editing in hESCs was performed as in ^53^. Guide RNAs (sgRNAs) were generated using the GeneArt Precision gRNA Synthesis Kit (Thermo Fisher Scientific). 0.6 μg sgRNA was incubated with 3 μg SpCas9 protein for 10 min at room temperature and electroporated into 2×10^5^ H9 cells using Neon transfection system (Thermo Fisher Scientific). Cells were put in a low O2 incubator and allowed to recover for 24-72 hours. Cells were then single cell sorted into 96-well plates with Sony Biotechnology (SH800S) Cell Sorter and grown up for 7-12 days. Individual clones were verified for out of frame deletions were verified via DNA sequencing with Illumina MiSeq and protein deletion was verified via immunoblotting. sgRNA target sequences were as follows: CCPG1 sgRNA TTCTAACTTAGGTGGCTCAA, TEX264 sgRNA CATGTCGGACCTGCTACTAC, FAM134A sgRNA TAATACGACTCACTATAG, FAM134B sgRNA GTCTGACACAGACGTCTCAG, FAM134C sgRNA AACTTGAGCTGTCAGACCAACA.

### Antibodies

The following antibodies were used:

ATG5 Rabbit Polyclonal Antibody Cell Signaling Technology 12994S, Lot5, WB 1:1000, RRID:AB_2630393; FAM134B Rabbit Polyclonal Antibody Proteintech 21537-1-AP, lot00100765, WB 1:1000 RRID:AB_2878879; FAM134C Rabbit Polyclonal Antibody Sigma-Aldrich HPA016492, lotR06641, WB 1:1000, RRID:AB_1853027; CCPG1 Rabbit Polyclonal Antibody Cell Signaling Technology 80158, lot1, WB 1:1000, RRID:AB_2935809; TEX264 Rabbit Polyclonal Antibody Sigma-Aldrich HPA017739, lot000012723, WB 1:1000, RRID:AB_1857910; REEP1 Rabbit Polyclonal Antibody Sigma-Aldrich HPA058061, lotR81573, WB 1:1000, RRID:AB_2683591; REEP4 Rabbit Polyclonal Antibody Sigma-Aldrich HPA042683, lotR39936, WB 1:1000, RRID:AB_2571730; REEP5 Rabbit Polyclonal Antibody Proteintech 14643-1-AP, lot00050540, WB 1:1000, RRID:AB_2178440; RTN3 Mouse Monoclonal Antibody Santa Cruz sc-374599, lot10922, WB 1:1000, RRID:AB_10986405; CKAP-4/p63 Sheep Polyclonal Antibody RD Biosciences AF7355, lotCGDGG012105B, WB 1:1000, RRID:AB_10972125; CKAP4 Rabbit Polyclonal Antibody Proteintech 16686-1-AP, lot0052093, WB 1:1000, RRID:AB_2276275; hFAB™ Rhodamine Anti-Tubulin Antibody BioRad 12004166, lot64512247, WB 1:10,000, RRID:AB_2884950; HSP90 mouse monoclonal Antibody Santa Cruz sc-69703, lotJ2721, WB 1:10,000, RRID:AB_2121191; GAPDH (D16H11) XP Rabbit Monoclonal Antibody Cell Signaling Technology 5174, lot8, WB 1:1000, RRID:AB_10622025; CREB-2/ATF-4 Mouse Monoclonal Antibody (B-3) sc-390063, lotJ2021, WB 1:1000, RRID:AB_2810998; VAPA Rabbit monoclonal Antibody Abcam ab181067, lotGR164232-2, WB 1:1000, RRID: AB_3073850; RTN1 (Isoform RTN1-C) Rabbit Polyclonal Antibody Proteintech 15048-1-AP, lot00043268, WB 1:1000, RRID:AB_2185981; Goat anti-Rabbit IgG HRP conjugate Bio-Rad 1706515, lot64559210, WB 1:3000, RRID:AB_11125142; Goat anti-Mouse IgG HRP conjugate Bio-Rad 1706516, lot64526160; WB 1:3000, RRID:AB_11125547; Neurofilament heavy polypeptide antibody Abcam ab7795, lotGR3448163-1, IF 1:300, RRID:AB_306084; MAP2 Guinea Pig Polyclonal Antibody Synaptic systems 188004, lot6-49, IF 1:300, RRID:AB_2138181; Nogo-A (C-4) (RTN4) Mouse Monoclonal Antibody Santa Cruz sc-271878, lotD2420, IF 1:300, RRID:AB_10709573; Calnexin Rabbit Polyclonal Antibody Proteintech 10427-2-AP, lot00094417, IF 1:300, RRID:AB_2069033; Goat anti-mouse Alexa488 Thermo Fisher Scientific A-11001, lot2379467, IF 1:300, RRID:AB_2534069; Goat anti-chicken Alexa488 Thermo Fisher Scientific A11039, lot218068, IF 1:300, RRID:AB_2534096; Goat anti-rabbit Alexa568 Thermo Fisher Scientific A-11011, lot2500544, IF 1:300, RRID:AB_143157; Goat anti-rabbit Alexa647 Thermo Fisher Scientific A27040, lot2659317, IF 1:300, RRID:AB_2536101; Goat anti-guinea pig Alexa488 Thermo Fisher Scientific A-11073, lot38320A, IF 1:300, RRID:AB_2534117; Goat anti-guinea pig Alexa647 Thermo Fisher Scientific A-21450, lot2446026, IF 1:300, RRID:AB_141882

### Western-Blotting

Cell pellets were resuspended in 8M Urea buffer (8M Urea, 150 mM TRIS pH, 150 mM NaCl) supplemented with protease and phosphatase inhibitor tablets and then sonicated twice, 10 seconds each, on ice. Lysates were clarified via centrifugation at 20,000 xg for 10 min at 4°C. BCA assays were performed on clarified lysates and normalized lysate amounts were boiled in 1X SDS containing Laemmeli buffer. Lysates were run on 4-20% Tris Glycine gels (BioRad) and transferred via Wet transfer onto PVDF membranes for immunoblotting with the indicated antibodies. Images of blots were acquired using Enhanced-Chemiluminescence or using the Rhodamine channel on a BioRad ChemiDoc imager and images quantified and converted to jpeg for publication using BioRad Image Lab Software RRID:SCR_014210.

### Flow Cytometry

hESCs that were converting to neurons were grown in 6-well plates and were treated with various drugs for the indicated time points and cell pellets were collected at the indicated day of neuronal differentiation. These were resuspended in FACS buffer (1X PBS, 2% FBS). At least 10,000 cells were analyzed on the Attune NxT (Thermo Fisher Scientific, Cat#A28993)) flow cytometer. Neutral Keima signal was measured at excitation at 445 nm and emission 603 nm with a 48 nm bandpass and acidic Keima signal was measured at 561 nm excitation and emission 620 nm and a 15 nm band pass. The resulting cell population Keima ratio was analyzed as previously described^54^. In brief, FCS files were exported into Flowjo where cells were gated for live cells, single cells and Keima positive cells. The 561(Acidic) to 445 (neutral) excitation ratio was calculated by dividing mean values of 561 nm excited cells to mean values of 445 nm excited cells.

### Imaging

Cells were plated onto 6 well, 12 well or 24 well glass bottom plates with high performance #1.5 cover glass (CellVis). Live cells were imaged at 37°C at 5% CO_2_. For immunofluorescence experiments, cells were fixed at room temperature with 4% paraformaldehyde plus in PBS, solubilized in 0.1% Triton-X in PBS and blocked with 1% BSA/0.1% Triton-X in PBS. Cell were then immunostained with anti-primary antibodies used at 1:500 and then AlexaFluor conjugated antibodies (Thermofisher) used at 1:300. Primary and secondary antibodies used in this study can be found in the Materials Table and described for each experiment detailed below. Fixed cell images were captured at room temperature. Cells were imaged using a Yokogawa CSU-X1 spinning disk confocal on a Nikon Ti-E inverted microscope at the Nikon Imaging Center in Harvard Medical School. Nikon Perfect Focus System was used to maintain cell focus over time. The microscope equipped with a Nikon Plan Apo 40x/1.30 N.A or 100x/1.40 N.A objective lens and 445nm (75mW), 488nm (100mW), 561nm (100mW) & 642nm (100mW) laser lines controlled by AOTF. All images were collected with a Hamamatsu ORCA-Fusion BT sCMOS (6.45 μm^2^ photodiode) with Nikon Elements image acquisition software.

#### Analysis of ER structures in axons

Cells were fixed and stained as described above specifically with α-Calnexin to detect ER, α-MAP2 to detect dendrites, α-NEFH to mark axons, and DAPI to detect nuclei. Z-stacks were acquired with the parameters stated above. Z series are displayed as maximum *z*-projections and brightness and contrast were adjusted for each image equally and then converted to rgb for publication using FiJi software. Fiji software was also used to split the z projections into individual channels for downstream image analysis in Cell Profiler^55^. Each field of view for all genetic backgrounds was thresholded in the same way with a consistent pipeline. The ‘identify primary objects’ tool was used to find nuclei, axons, dendrites, and ER structures. The α-NEFH-positive axon object regions were used to create an axon mask and ER structures within this mask were counted. The area of each ER structure was also measured. The number of ER axonal structures was then compared to the number of detected nuclei.

#### Analysis of cell nuclei using size

We first assayed if nuclei were in tact in the images used to assess the amount and size of ER structures in axons as described above (α-Calnexin to detect ER, α-MAP2 to detect dendrites, α-NEFH to mark axons, and DAPI to detect nuclei and z projections already split into individual channels as detailed above for downstream image analysis in Cell Profiler^55^). The DAPI channel images for all genetic backgrounds was thresholded in the same way with the following pipeline: two different ‘identify primary objects’ modules were used to find and count nuclei structures. In one, only larger “intact” nuclei were selected and counted (as was done previously for the analysis of ER structures in axons to get ER structures per nuclei). In the second, smaller fragmented nuclei were included in the thresholding method. The ratio of intact to total DAPI-positive nuclei structures was calculated and reported for each condition.

#### Analysis of cell nuclei using TUNEL

As secondary confirmation that the intact nuclei that we were assaying were indeed healthy, we performed a Click-iT Plus TUNEL (terminal deoxynucleotidyl transferase dUTP nick end labeling) assay (Invitrogen C10617, Alexa Fluor 488), which detects DNA breaks formed when DNA fragmentation occurs at the end of apoptosis. We prepared four new differentiations of WT, ATG12 and PKO neurons at day 20 to perform this staining. In short, following the kit protocol, after fixing and permeabilizing the iNeurons as described above we followed the kit directions to first perform a TdT reaction. In this reaction, the TdT enzyme takes EdUTP (a dUTP modified with a small, bio-orthogonal alkyne moiety) and incorporates it at the 3’-OH ends of fragmented DNA. Next, we performed the click reaction, a copper catalyzed covalent reaction occurs between the Alexa Fluor™ picolyl azide dye and an alkyne. Furthermore, detection of the DNA break is based on Alexa Fluor signal at that site. After performing this Click-iT Plus TUNEL reaction, we next stained with DAPI to label all DNA structures (labels both intact and fragmented DNA). Z-stacks were acquired with the parameters stated above. Z series are displayed as maximum *z*-projections and brightness, and contrast were adjusted for each image equally and then converted to rgb for publication using FiJi software. Fiji software was also used to split the z projections into individual channels for downstream image analysis in Cell Profiler^55^. For the TUNEL channel images, images from all genetic backgrounds were thresholded in the same way using an ‘identify primary objects’ modules to find and count all damaged DNA structures, including larger and smaller structures. For the DAPI channel images, two different ‘identify primary objects’ modules were used to find and count DAPI structures. In one, only larger “intact” DAPI-positive nuclei were selected. In the second, smaller fragmented DAPI-positive nuclei were included in the thresholding method. To calculate the total nuclei number, the number of damaged TUNEL-positive DNA structures was added to the number of intact DAPI nuclei. In the final analysis, the ratio of intact DAPI-positive nuclei structures to total nuclei (damaged TUNEL positive nuclei plus intact) was calculated and reported for each condition.

#### Visualizing Keima-ER in neuronal differentiation

Live cells stably expressing Keima-RAMP4 (localizes to all ER) or Keima-REEP5 (localizes to ER tubules specifically) were imaged at the indicated day in neuronal differentiation. Pairs of images for ratiometric imaging of Keima-RAMP4 fluorescence were collected sequentially using 100 mW 442 nm (neutral Keima excitation) and 100 mW 561 nm (acidic Keima excitation) solid state lasers and emission collected with a 620/60 nm filter (Chroma Technologies). Z-stacks were acquired with a Nikon Plan Apo 40×/1.45 N.A oil-objective lens. Z series are displayed as maximum z-projections and brightness and contrast were adjusted for each image equally and then converted to rgb for publication using FiJi software. Fiji software was also used to split the z projections into individual channels. For each channel, complimentary line scans 30 μm long with 1.7 μm width were drawn in either the soma or projection of iNeurons. The 561 nm or 442 nm gray values along these lines was measured using ‘plot profile’ in Fiji. The 561/442 ratio of these values at each complimentary point along the line was calculated and plotted in excel.

#### Characterizing spatial and temporal properties of ER-phagy receptors

hESCs with WT or ATG12^-/-^ genetic background stably expressing WT or mutant TEX264-GFP or FAM134C-GFP were converted to neurons and treated with various drugs for the indicated time points and imaged at the indicated day in neuronal differentiation. Z-stacks were acquired with the parameters stated above. Z series are displayed as maximum *z*-projections and brightness and contrast were adjusted for each image equally and then converted to rgb for publication using FiJi software.

For day 4 cells (untreated or treated with the indicated drugs), the number of GFP puncta per cell was quantified using Cell Profiler. Each field of view for all genetic backgrounds and drug treatments was thresholded in the same way with a consistent pipeline. Using the ER-phagy receptor (488ex, GFP channel) max z projection image, the ‘identify primary objects’ tool was used to detect cells (receptor labels the whole ER membrane which can be used to identify the cells) and to detect puncta (small bright circles found within the ER membrane). The puncta were linked to each cell and the puncta per cell numbers were exported.

Autophagosome (LC3B) and ER-phagy receptor (TEX264 or FAM134C) co-labelling was achieved by transducing with mCh-LC3B and receptor-GFP lentivirus. Day 30 neurons were imaged live for 30 min with an image acquired every 30 sec. Fiji was used to track GFP and mCh positive puncta. Lines between each frame were used to measure the distance traveled of the puncta from frame to frame. Forward direction was reported as a positive value in micron and backward direction was reported as a negative value. Events in neurons from three independent differentiations were captured. The events were binned based on their speed of movement in micron per second. The percentage of events at each speed were plotted as using GraphPad Prism 7.

After live cell imaging at day 30, the ER-phagy receptor and mch-LC3B positive transduced neurons were fixed as described above. The iNeurons were immunostained with α-MAP2 to detect dendrites and α-NEFH to mark axons. Z-stacks were acquired with the parameters stated above. Z series are displayed as maximum *z*-projections and brightness and contrast were adjusted for each image equally and then converted to rgb for publication using FiJi software.

### RNA extraction, Reverse transcription-PCR, DNA gel electrophoresis

At day 12, iNeurons with each genotype was left untreated or was treated with tunicamycin. At 4hr, all the cells were scrapped off the dishes, pelleted and washed with three times with PBS. Number of cells was determined and then the pellets were snap frozen in liquid nitrogen and stored at -80C for a few days before use. Cell pellets were thawed and resuspended in freshly prepared RLT buffer (350 μL per sample for 1E6 cells) from the RNAeasy Qiagen kit (Qiagen 74104). Dnase1 digestion buffer was then added and cells were subsequently lysed via passage through a Qiashredder column (Qiagen 79654). One volume of 70% ethanol was added to the lysate and the lysate-EtOH solution was transferred to a RNAeasy spin column. The following spins, on column DNAseI (Thermo EN0521) digestion, buffer washes, and RNA elution were performed via the RNAeasy Qiagen kit directions. Final extracted RNA concentration for each condition was measured using a anorop. Reverse transcription reactions for each condition (using the same amount of starting μg of RNA, 0.5 μg, in each reaction) were performed with Superscript III reverse transcriptase master mix (Invitrogen 18080-051) using oligo dT_20_ primers (Invitrogen 79654) and dNTPs (NEB N0447L) to create complementary DNA (cDNA). With the cDNA, PCR reactions were performed to amplify cDNA from GAPDH mRNA (forward 5′GGATGATGTTCTGGAGAGCC3′, reverse 5′CATCACCATCTTCCAGGAGC3′) ; or to amplify cDNA from unspliced XBP1 mRNA or spliced XBP1 mRNA (forward 5′ CCTTGTAGTTGAGAACCAGG 3′, reverse 5′GGGGCTTGGTATATATGTGG 3′) (as performed in ^56,57^. PCR products were electrophoresed on a 2.5 percent agarose gel. The size difference between the spliced and the unspliced XBP1 is 26 nucleotides.

### Transmission Electron Microscopy

#### Cell preparation

iNeurons were grown on aclar plastic discs in 12-well plates coated with Matrigel. At day 20, the iNeurons were fixed with 2.5% Glutaraldehyde 1.25% Paraformaldehyde and 0.03% picric acid in 0.1 M sodium cacodylate buffer (pH 7.4). A 2x solution was diluted 1:1 with the cell media in the dish. Cells were fixed at room temperature for one hour.

#### Epon Embedding

The following steps were performed by the Harvard Medical School Electron Microscopy Core: Cells were washed in 0.1 M sodium cacodylate buffer (pH 7.4), postfixed for 30 min in 1% Osmium tetroxide (OsO4)/1.5% Potassiumferrocyanide (KFeCN6), washed in 2x in water and 1x in Maleate buffer and incubated in 1% uranyl acetate in Maleate Buffer for 30min followed by 2 washes in water and subsequent dehydration in grades of alcohol (5min each; 50%, 70%, 95%, 2x 100%). The samples were subsequently embedded in TAAB Epon (TAAB Laboratories Equipment Ltd, https://taab.co.uk) and polymerized at 60C for 48 hrs. Note on Embedding –a drop of Epon is put onto a clean piece of Aclar, then the coverslip is removed from 100% EtOH with a pair of fine tipped forceps. Excess EtOH is removed by quickly blotting the side of the coverslip onto a filter paper (to make sure the cells don’t dry out) and the coverslips are placed (cell side down) onto the Epon drop. A small weight on top helped with keeping it flat. After polymerization, the Aclar was peeled off, a small area (∼1mm) of the flat embedded cells was cut out with a razor blade and remounted on an Epon block. Ultrathin sections (about 80nm) were cut on a Reichert Ultracut-S microtome, picked up on to copper grids stained with lead citrate and examined in a TecnaiG^2^ Spirit BioTWIN and images were recorded with an AMT 2k CCD camera. Regions close to the coverslip were specifically targeted to capture neuronal processes, not somata.

### Cryo-Electron Tomography

#### Cryo-ET sample preparation and freezing

AAVS-TREG3-NGN2 non-embryonic and internationally accepted iPSCs were differentiated to iNeurons and cultured on EM grids as described in detail in this protocol: https://www.protocols.io/view/neural-differentiation-on-em-grids-ineurons-sample-5jyl8jz36g2w/v2. In this particular case, a 200-mesh gold grid with Silicon Dioxide R2/1 film (Quantifoil) was coated with Matrigel as reported in the protocol. The iPSC-derived iNeurons on the grid were transduced at Day 12 with Lentiviruses carrying mCherry-LC3B and TEX264-GFP. Medium was gradually and completely exchanged in the next days with ND2 without phenol red (prepared using phenol red-free Neurobasal, Thermo Fisher, Gibco 12348017). iNeurons were plunged on DIV 18 with a Vitrobot Mark IV (Thermo Fisher Scientific), with application of 4 μl of phenol red-free ND2 medium and with the following settings: room temperature, humidifier 70%, Blot-Force 8, Blot-Time 9 s.

#### Cryo-Fluorescence data acquisition and processing

Fluorescence stacks of grid squares of interest were acquired before tilt-series acquisition on a SP8 cryo-confocal laser scanning microscope equipped with a cryo stage and a 0.9 NA/50X objective (Leica). Stacks were acquired sequentially in the red (ex. 552 nm / em. 598-625 nm) and green channel (ex. 488 nm / em. 498-525 nm) using hybrid detectors, with an x/y pixel size of 60 nm and a z step size of 400 nm. Transmitted light data were collected simultaneously to visualize positions of the support film holes for 2D registration and correlation. Fluorescence data were deconvolved with Huygens (SVI, The Netherlands) using a theoretical PSF and CMLE method. Maximum intensity projections (MIPs) of the fluorescence channels stacks were generated in Fiji^58^. For transmitted light data, one slice focused on the holes of the support film was selected and then used for the 2D correlation procedure.

#### Cryo-ET data acquisition and Processing

TEM data acquisition was performed on a Krios G4 at 300 kV with Selectris X energy filter and Falcon 4i camera (Thermo Fisher Scientific, Eindhoven, The Netherlands) using Tomo5 (Version 5.12.0, Thermo Fisher Scientific). Tilt series were acquired at a nominal magnification of 42,000X (pixel size 2.93 Å) using a dose-symmetric tilt scheme with an angular increment of 2°, a dose of 2 e^-^/Å^2^ per tilt and a target defocus between -2.5 and -4 μm. Tilt series were collected ranging from -60° to 60° with a total dose of 120 e^-^/Å^2^, and frames were saved in the EER file format. The positions for tilt series acquisition were determined by visual inspection of 11500X magnification “search” montage maps acquired in thin areas of the sample (**Extended Data Fig. 5a**). Tilt series were recorded where double membrane vesicle structures could be seen inside intact iNeuron projections, with continuous plasma membrane and microtubules bundles. Most of the autophagosomes were captured in areas in which the sample was thicker than 400 nm (**Extended Data Fig. 5o**). While such a high sample thickness is generally not recommended for subtomogram averaging of particles because of the low SNR of the resulting images, it still allowed neural-network based segmentation and visualization of autophagosomes and their membrane cargo. Tilt series frames were motion-corrected with Relion’s implementation of Motioncorr2 for EER files ^59^ and reconstruction was performed in IMOD (v.4.10.49, RRID:SCR_003297, https://bio3d.colorado.edu/imod/) by using the TomoMAN wrapper scripts https://doi.org/10.5281/ZENODO.4110737. Tomograms at 2× binning (IMOD bin 4) with a nominal pixel size of 1.172 nm were denoised using cryo-CARE ^60^ (https://github.com/juglab/cryoCARE_T2T).

#### Cryo-ET dataset annotation and analysis

All double membrane compartments with (1) relatively tight and regular intermembrane spacing and (2) near-spherical shape were identified as autophagosomes. Note that while some autophagosomes presented a tight and homogeneous intermembrane distance (**Extended Data Fig. 5n**, autophagosomes 3 and 4), others presented irregularities in the distance, or small bumps in the inner membrane (**Fig. 4f**), which might suggest that they are amphisomes – autophagosomes that have already fused with one or few small lysosomes^61^. N=32 tomograms contained autophagosomes and some of them contained more than one, leading to total count of n=37 autophagosomes. All structures were first annotated by cargo: Tubular membrane cargo structures, morphologically similar to the ER tubules present in the cytoplasm of the same tomograms (see **Fig. 4c, Extended Data Fig. 5e**), were labeled as “tubular ER cargo”. Note that autophagosomes often contained single membrane vesicles cargo next to tubular ER (**Fig. 4c,f and Extended Data Fig. 5n**). Second, autophagosomes were annotated as “microtubule-linked’ when microtubules were found at a distance closer than 20 nm to their outer membrane (**Extended Data Fig. 5f**). Tomogram thickness was determined in 3Dmod (IMOD) by measuring the distance between the upper and lower boundary of the sample, where small pieces of ice contamination are often visible (**Extended Data Fig. 5o**). Plots were generated with Python 3.9.7 using pandas 1.3.0 (https://pandas.pydata.org/, RRID:SCR_018214) ^62^, matplotlib 3.3.0 (https://matplotlib.org/, RRID:SCR_008624) ^63^ and seaborn 0.11.0 (https://seaborn.pydata.org/, RRID:SCR_018132)^64^ packages.

#### 2D correlation of cryo-fluorescence data on TEM images

Correlation of all autophagosomes with the previously acquired cryo-fluorescence data was investigated through a two-step procedure using Fiji’s BigWarp plugin^65^ as described in the following. First, centers of different holes on the support film are selected both in an 800X TEM “overview” image and in the transmitted light image previously acquired on the cryo-confocal microscope (**Extended data Fig. 5g**). After registration, the 800X TEM image is transformed (affine transformation) and overlayed on the green and red fluorescence MIPs (**Extended Data Fig. 5h**). The overlay is then cropped to obtain a subregion around the tomogram position which is used for the second correlation with the 11500X TEM “search” image (**Extended data Fig. 5i**). This time, fine landmarks such as pieces of ice, features of the cellular sample and positions along the hole are selected and used for transforming the cropped fluorescence data (affine transformation). A final overlay of the 11500X TEM search image with the cryo-fluorescence data is generated to visualize correlation of autophagosomes with fluorescence signals (**Extended Data Fig. 5k**,**m**). This procedure yielded a total of n=5 out of 37 autophagosomes coinciding with distinct TEX264-GFP signal peaks (**Fig. 4b)**. All the 5 tomograms coinciding with TEX264 contained tubular ER cargo, and 4 of them were in close proximity to microtubules (**Extended Data Fig. 5l**). Note that the LC3B cryo-fluorescence signal often appears diffuse and bright in the cytosol, and it was difficult to distinguish peaks of signal in correspondence of autophagic structures (**Extended Data Fig. 5k**,**m**). This might be due to LC3B localizing also on microtubules or distributed throughout the cytosol and its involvement in other cellular processes, such as non-canonical autophagy^35^ and LC3-associated phagocytosis ^66^. Consequently, LC3B-mCherry signal could not be used for reliable cryo-correlation in this case. Moreover, often extracellular debris, pieces of ice and other intracellular structures can be autofluorescent at cryogenic temperature in the green and/or red channel. This phenomenon has been previously reported by others and increases the noise in the cryo-fluorescence images^67^.

#### Tomogram segmentation

Segmentations were done for the 5 tomograms of autophagosomes coinciding with TEX264 signal. Membranes were segmented with Membrain-Seg (https://github.com/teamtomo/membrain-seg), a U-Net based tool for membrane segmentation in cryo-ET data, using the publicly available “best” pretrained model (v9). This method reliably detected membranes, also in very thick tomograms (**Extended Data Fig. 5n**,**o**). However, it often merged membranes corresponding to different compartments and sometimes picked cytoskeletal components such as microtubules and neurofilaments in the very crowded neuronal subcellular environment. To separate the different membrane compartments, a watershed segmentation was performed on the original Membrain-seg output, using as seeds a “one-click” rough segmentation of different membranes generated in Amira (Thermo Fisher Scientific) from the output of tomosegmemTV^68^. For thicker tomograms, such as autophagosome 3 and 4 (**Extended Data Fig. 5n**,**o**), automated segmentations of the cargo ER were manually refined in Amira. For autophagosomes 1 and 2 (**Fig. 4c**,**f**), microtubules were segmented automatically with Dragonfly (version 2022.2, Comet Technologies Canada Inc., Montreal, Canada.). A custom model was trained for each of the tomograms, following a previously described protocol^69^. 3D renderings of the segmentations were generated in ChimeraX^70^.

### Quantitative proteomics

#### Sample preparation for Mass Spectrometry

Cell pellets were resuspended in 8M Urea buffer (8M Urea, 150 mM TRIS pH, 150mM NaCl) supplemented with protease and phosphatase inhibitor tablets and then sonicated twice, 10 seconds each, on ice. Lysates were clarified via centrifugation at 20,000 xg for 10 min at 4°C. BCA assays were performed on clarified lysates. 100 ug of each sample was taken and total volume raised to 100 μL total. Samples were reduced using TCEP (0.5 M for 30 min at room temperature) and alkylated (with Chloroacetamide (20 mM for 20 min at room temperature) prior to methanol-chloroform precipitation with 3:1 methanol, 1:1 chloroform, and 2.5:1 water added. Aqueous and organic phases were separated with centrifugation for 5 min at 14,000 xg. Liquid around the protein layer was removed and this protein layer was washed with 1 mL methanol and then pelleted at 5 min at 14,000 xg. The supernatant was removed. The pellets were then resuspended in in 50μL, 200 mM EPPS, pH8.5. Peptide digestion was carried out using LysC (1:100) for 2h at 37°C followed by Trypsin (1:100) overnight. 25 μL of the digested peptides were then labelled by adding 5 μL 100% acetonitrile (CAN) and with 7 μL of TMT reagent (at 20 mg/ml stock in ACN) for 2h and the reaction was quenched using hydroxylamine at a final concentration of 0.5% (w/v) for 20 min.

#### Basic pH reversed-phase

Samples were combined 1:1 such that each channel consisted of the same amount of peptide. The pooled peptide sample was desalted with a 100 mg Sep-Pak solid phase extraction column and then fractionated with basic pH reversed-phase (BPRP) HPLC. Fractionation was executed using an Agilent 1200 pump with an Agilent 300 Extend C18 column (3.5 μm particles, 2.1 mm ID, and 250 mm in length). A 50 min linear gradient from 5% to 35% acetonitrile in 10 mM ammonium bicarbonate pH 8 at a column flow rate of 0.25 mL/min was used for peptide fractionation. A total of 96 fractions were collected and then concatenated down to 24 superfractions, as described previously ^71^. These 24 superfractions were divided into two sets of 12 non-adjacent superfractions and were acidified by adding formic acid to a concentration of 1%. One set of fractions (n=12) were vacuum centrifuged to near dryness, and each was desalted via StageTip, dried by vacuum centrifugation, and reconstituted in 5% acetonitrile, 5% formic acid prior to LC-MS/MS analysis.

#### Mass spectrometry data acquisition and processing

Mass spectrometric data were collected on an Orbitrap Fusion Lumos mass spectrometer coupled to a Proxeon NanoLC-1200 UHPLC and a FAIMSpro interface ^72^. The 100 μm capillary column was pulled in-lab and packed with 35 cm of Accucore 150 resin (2.6 μm, 150 Å; ThermoFisher Scientific). Peptides were eluted over a gradient of 90 or 110 min. consisting of 5% acetonitrile to 30% acetonitrile in 0.125% formic acid. The scan sequence began with an MS1 spectrum (Orbitrap analysis, resolution 60,000, scan range 350–1350 or 400-1600 Th, automatic gain control (AGC) target was set as “standard,” maximum injection time was set to auto). SPS-MS3 analysis was used to reduce ion interference ^73,74^. MS2 analysis consisted of collision-induced dissociation (CID) and quadrupole ion trap analysis (automatic gain control (AGC) 2 ×10^4^, normalized collision energy (NCE) 35, q-value 0.25, maximum injection time 35 ms, isolation window 0.7 Th). Following the acquisition of each MS2 spectrum, we collected an MS3 spectrum in which multiple MS2 fragment ions were captured in the MS3 precursor population using an isolation waveform with multiple frequency notches. MS3 precursors were fragmented by higher-energy collisional dissociation (HCD) and analyzed using the Orbitrap (NCE 55, AGC 1.5 ×10^5^, maximum injection time 150 ms, resolution 50,000). We used the Real Time Search (RTS) using Orbiter^75^ with a *H. sapiens* database (UniProt, downloaded August 2020) and we limited MS3 scans to 2 peptides per protein per fraction. A total of 24 RAW files were collected with data for 12 non-adjacent superfractions acquired using a compensation voltage (CV) set of -40/-60/-80V with a 1.25-sec TopSpeed cycle was used for each CV.

Spectra were converted to mzXML via MSconvert ^76^. Database searching included all *H. sapiens* entries from UniProt. The database was concatenated with one composed of all protein sequences in that database in the reversed order. Searches were performed using a 50-ppm precursor ion tolerance for total protein level profiling. The product ion tolerance was set to 0.9 Da. These wide mass tolerance windows were selected to maximize sensitivity in conjunction with SEQUEST ^77^ searches and linear discriminant analysis ^78,79^. TMT labels on lysine residues and peptide N termini (+304.207 Da) and carbamidomethylation of cysteine residues (+57.021 Da) were set as static modifications, while oxidation of methionine residues (+15.995 Da) was set as a variable modification. Peptide-spectrum matches (PSMs) were adjusted to a 2% false discovery rate (FDR) ^80,81^. PSM filtering was performed using a linear discriminant analysis, also as described previously ^79^ and then assembled further to a final protein-level FDR of 2% ^81^.

Proteomics Data analysis. Peptide-spectral matches (PSM) were filtered for summed signal-to-noise ratio (SNR>200) across the TMT plex and for precursor signals that contained > 0.5 isolation purity of the MS1 isolation window. To normalize protein input across TMT channels, all PSM intensities were summed and the total intensity per channel were sum normalized to the median summed intensity across the TMTpro plex. Protein intensities were generated by summing input-normalized TMT intensities for its constituent peptides’ PSMs^82^, serving as a weighted average quantification. Comparison among experimental conditions (n=3-4 biological replicates) were conducted by performing a Student’s t-test of normalized log_2_-transformed protein TMT intensities. Resulting p-values were adjusted for multiple hypothesis correction using the Benjamini-Hochberg approach^83^. For heatmap generation or linear model analysis, replicate protein report ion intensities were normalized to the mean of the biological replicates of either day0 for the differentiation experiment or to wildtype control day12 iNeurons replicates within a given TMTpro plex.

To conduct the linear regression analysis using a single model for the additive combinatorial ER receptor knockout TMT data, we incorporated indicators/dummy variables that can take on one of two possible numerical values (1: contains addition of an ER receptor knockout(s) or 0: does not). All replicates were normalized to mean of WT control which was centered at 0, essentially removing the intercept estimation (β_0_) from the model. This was because the TMT protein reporter intensities are not indicative of absolute abundance, and we are interested in understanding the fold change contribution from the addition of each ER receptor knockout. The following indicators/dummy variables were used:

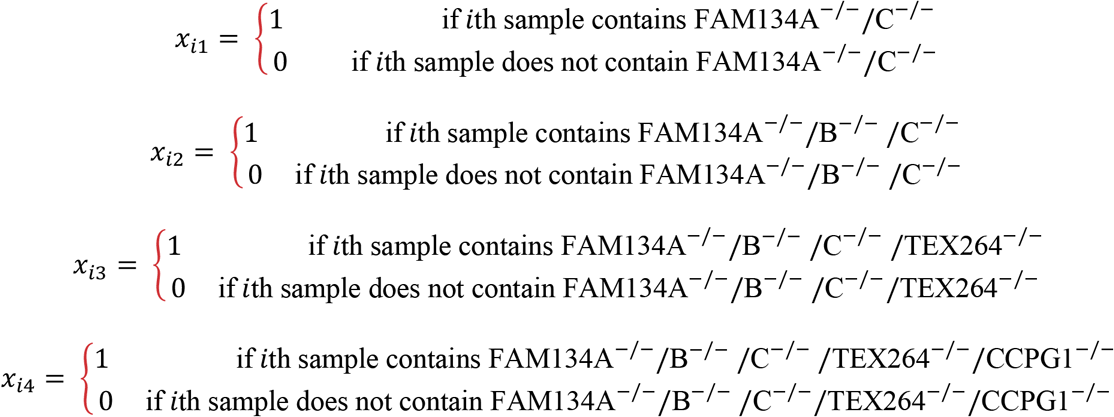

The model below was then used to estimate betas (β) for each step-wise addition of ER receptor knockout(s) with inherent technical noise from the MS acquisition and reporter quantification.

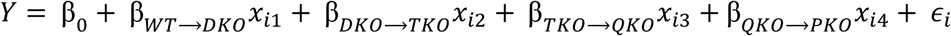

Thus:

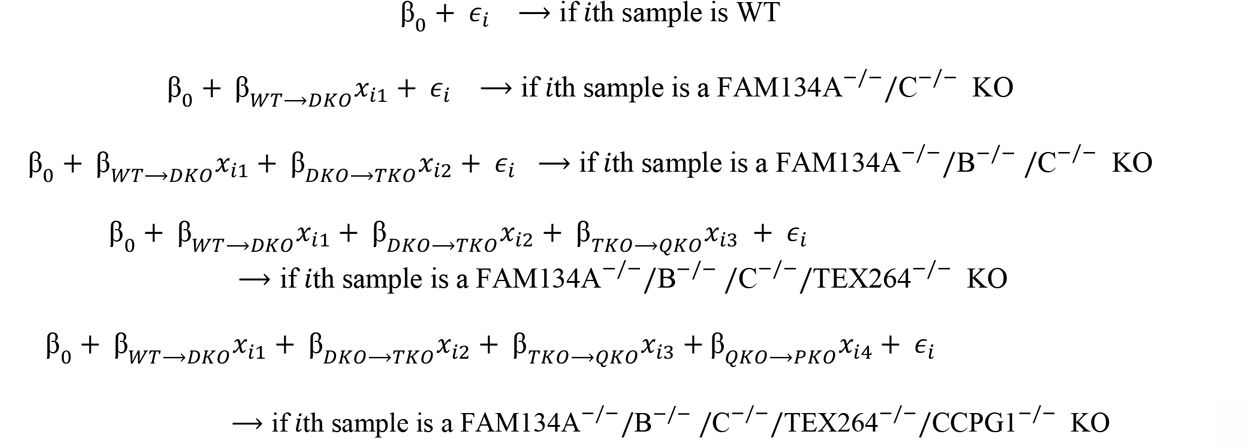

In R using the *lm* function, the beta (β) coefficients and p-values were extracted from the model, and beta (β) coefficients and Benajmini-Hochberg^83^ adjusted p-values were leveraged for downstream analysis and figure generation. One can interpret the β_TKO→QKO_ for instance as the average foldchange from the triple knockout to the quadruple knockout, due to the addition of TEX264 KO on the FAM134A^-/-^/B^-/-^/C^-/-^ knockout cells.

Classification of proteins to various organellar locations or functional groups were performed using manually curated databases from Uniprot and are listed in the relevant supplementary tables. Sub-cellular annotations were derived from Itzak et al.^27^ with additional cytosol protein location designations from Uniprot. ER high sheet and high curvature annotations were extracted from ref^2^.

### Proteomics Data Availability

The mass spectrometry proteomics data have been deposited to the ProteomeXchange Consortium via the PRIDEpartner repository^84^ with the dataset identifiers PXD041069 and PXD046646 and reviewers can access with username..

### Code Availability

Code for proteomics data analysis and relevant figure generation can be found on GitHub at the https://github.com/harperlaboratory/iNeuron_ERphagy.git repository.

### Statistics

Proteomics data analysis was performed using R (4.2.2) within the Rstudio IDE (2022.12.0 Build 353, Posit). Data visualizations in the form of heatmaps, volcano plots, violin plots, protein abundance profiles, and subcellular localization plots were generated using the following R packages: tidyverse (2.0.0), dplyr (1.0.10), cowplot (1.1.1), pheatmap (1.0.12), stringr (1.5.0), RColorBrewer (1.1-3), ggrepel (0.9.2), ggplot2 (3.4.1), purr (1.0.1), and tibble (3.1.8). For imaging statistics, GraphPad Prism9 was used. Mean (for number of ER structures per nuclei), mean (for the area of axonal ER structures), percent intact nuclei, and percent TUNEL negative nuclei values from each replicate differentiation experiment (n=4 in each experiment) were compared between each knockout and wildtype using a Mann-Whitney test. For flow cytometry quantification, GraphPad Prism9 was used. Each condition had three biological replicates. Brown-Forsythe and Welch One-way ANOVA and Dunnett’s T3 multiple comparisons test (assuming a Gaussian distribution) were used to compare each condition. For imaging and flow cytometry analysis *, p<0.05; **, p<0.01, ***, p<0.001. For proteomics datasets, the alpha used for FDR cut-offs was adjusted p-value <0.05 to consider significance. To compare log_2_FCs for each organelle proteome to a random distribution, a randomized protein selection was generated (100 iterations) keeping the same number of proteins as the perspective organelle. The log_2_FC distribution of this random protein set was compared to the organelle log_2_FC distribution using a Kolmogorov-Smirnov Test (two-sided) test. For analysis of violins, when calculating the degree of change of each ER compartment’s β-values from no change (zero), a Wilcoxon one sided test was used with Bonferroni p-value correction applied due to the multiple comparisons. For other analysis of violins comparing the log2FCs between two genotypes for each ER compartment, the comparison was made using paired Wilcoxon two sided tests. All data figures were generated in Adobe Illustrator using R (4.1.3), Rstudio IDE(2021.09.3 Build 396, Posit), and GraphPad Prism9. Unless stated otherwise, all quantitative experiments were performed in triplicate and average with S.E.M. or S.D. as indicated in legends reported.

